# NRF2 upregulation by CDDO-Me protects AC16 human cardiomyocytes against doxorubicin-induced toxicity

**DOI:** 10.1101/2025.09.10.675420

**Authors:** James A. Roberts, Michael Batie, Amy H. Ponsford, Jonathan Poh, Benjamin J. Hewitt, Hannah F. Botfield, Lisa J. Hill, Christopher M. Sanderson, Sonia Rocha, Parveen Sharma

**Author notes:** Robert Aitken Institute for Clinical Research, University of Birmingham, Birmingham, UK, B15 2TT., Sherrington Building, University of Liverpool, Liverpool, UK, L69 3GE.

## Abstract

Doxorubicin (DOX) is an effective anticancer therapeutic but exhibits dose-dependent, potentially life-threatening cardiotoxicity. The specific mechanisms driving this cardiotoxicity are not fully understood but can include the induction of oxidative stress and subsequent cell death mechanism activation. This has prompted the exploration of NRF2, a master co-ordinator of antioxidant and largely cytoprotective pathways, as a potential approach for the alleviation of DOX-induced cardiotoxicity. Here, NRF2 was pharmacologically activated via CDDO-Me (hitherto referred to as CDDO) to reduce the negative consequences on AC16 human cardiomyocyte cell health and functions. NRF2 intracellular dynamics were quantitatively measured using live-cell imaging, demonstrating rapid (∼10 min) yet sustained (≥24 h) induction of NRF2 expression and functional downstream activity. Genetic perturbations of the NRF2-KEAP1 system highlight that CDDO acts specifically through NRF2 to exert AC16 cytoprotection from DOX whilst not promoting human lung and pancreatic cancer cell line viability. Via RNA-seq analysis, we reveal that CDDO dampens DOX-mediated effects on p53 signalling, apoptosis and ferroptosis. This study provides novel insight into NRF2 dynamics in the widely utilised AC16 cells whilst further elucidating the molecular mechanisms contributing to DOX cardiotoxicity and potential NRF2-orchestrated defence.

**Graphical abstract:** 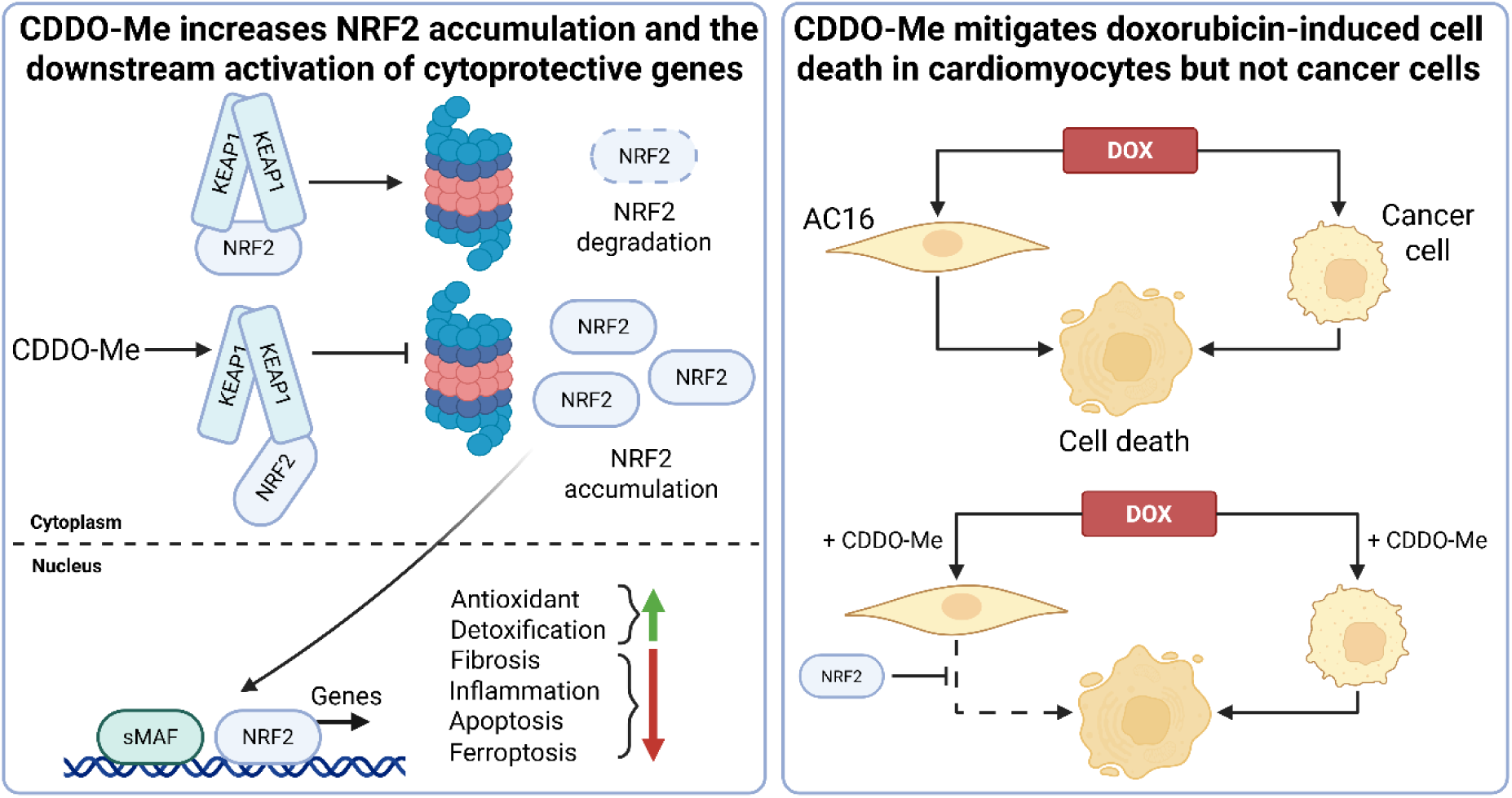

## 1 Introduction

Doxorubicin (DOX) is an anthracycline chemotherapeutic which has been utilised since the 1960s to target solid and haematological cancers [1,2] including ∼60% of childhood malignancies [3]. However, the clinical use of DOX is constrained due to its harmful adverse effects on healthy tissues, most prominently of the heart [4]. DOX exhibits cumulative, dose-dependent, irreversible and life-threatening cardiotoxicity which may present years or decades following treatment cessation [5]; clinical manifestations include reduced left-ventricular ejection fraction, myocardial strain, arrhythmia, cardiomyopathy and heart failure [6–8]. Despite the longstanding utility, the biochemical mechanisms contributing to DOX-induced cardiotoxicity are not fully elucidated due to its multifaceted mechanistic nature. DOX intercalates with DNA, inhibits topoisomerase II (TOP2), triggers the induction of double-stranded breaks and prevents repair and replication mechanisms [1,9]. DOX also triggers oxidative stress which, in turn, largely contributes to cell toxicity via the activation of cell death mechanisms, leading to cardiac tissue dysfunction [10–13]. The heart is particularly vulnerable to oxidative stress due to low basal expression of antioxidant enzymes, high abundance of the reactive oxygen species (ROS)-producing mitochondria and inability to replenish damaged cardiomyocytes due to their quiescence in a mature state [14–16].

NF-E2 p45-related factor 2 (NRF2), encoded by the *NFE2L2* gene, is a cap’n’collar basic-region leucine zipper transcription factor considered as a master regulator of antioxidant responses and thus has emerged as a therapeutic target to alleviate DOX-induced cardiotoxicity [8,17]. Under basal conditions, NRF2 activity is restrained predominantly through binding to KEAP1 homodimers which facilitate the recruitment of ubiquitin ligases and polyubiquitination of the NRF2 protein, ultimately priming NRF2 for proteasomal degradation and constitutive repression [18,19]. Thus, the NRF2 half-life may be as short as 10 min [20]. However, upon increasing ROS or electrophile abundance, thiol-rich cysteines within KEAP1 are chemically modified which prevents the ubiquitination of NRF2 and permits the nuclear translocation of *de novo* NRF2 molecules [21,22]. Here, NRF2 heterodimerises with small musculoaponeurotic fibrosarcoma (sMAF) proteins and binds to targets within the DNA harbouring antioxidant/electrophile responsive elements (ARE/EpRE) to upregulate genes associated with antioxidation, iron regulation, bioenergetics, and autophagy amongst other functions [22–24].

Pharmacological upregulation of NRF2 activity has previously shown a protective phenotype against DOX cardiotoxicity *in vitro* [25,26] and *in vivo* [27]. Despite this, the specific mechanisms driving such cardiovascular-preserving outcomes are yet to be fully explored and may extend beyond the antioxidant activity of NRF2. Here, the impact of 2-cyano-3,12-dioxo-oleana-1,9(11)-dien-28-oic acid methyl ester (CDDO-Me), herein referred to as CDDO, a potentially specific post-transcriptional activator of NRF2 via modulation of a single KEAP1 cysteine residue (Cys151) [28], was utilised to illuminate the biomolecular processes resulting in the alleviation of DOX-induced toxicity in human cardiomyocytes. Moreover, given the propensity for therapeutic resistance in tumours with aberrant upregulation of NRF2 [29–31] and the cancer-targeting clinical utility of DOX, cardio-protective phenotypes were explored in human cancer cells. To better elucidate the specific mechanisms contributing to DOX cardiotoxicity and the potential protection afforded via NRF2 stimulation, RNA-seq analysis was concurrently performed. Excitingly, this research strongly suggests that CDDO promotes cardiomyocyte protection against DOX cytotoxicity directly in an NRF2-dependent manner. Importantly, CDDO does not promote the survival of multiple cancer types in the presence of DOX. As such, CDDO presents as a candidate molecule for the alleviation of cardiotoxicity in patients receiving DOX and other chemotherapeutics with adverse cardiovascular events.

## 2 Materials and methods

### 2.1 Materials

All cell culture and chemical reagents were purchased from Invitrogen, USA or Sigma, USA, respectively, unless otherwise stated. DOX was purchased as DOX HCl from Selleckchem, USA.

### 2.2 Cell culture

All cell lines were cultured in Dulbecco’s Modified Eagle’s Medium (DMEM) with GlutaMAX, 25 mM glucose, 1 mM pyruvate and 1% non-essential amino acids (NEAA) with supplemented 10% fetal bovine serum (FBS), 100 U/mL penicillin and 100 µg/mL streptomycin antibiotics. Cells were grown within a humified atmosphere at 37°C and 5% CO_2_ and maintained through regular passaging; experiments were performed on cells between passages 3-10. AC16 human cardiomyocytes and A549 human lung adenocarcinoma parent cell lines were obtained from ATCC, USA. SUIT-2 and MIA PaCa2 human pancreatic cancer cell lines were kindly provided by the Costello-Goldring lab, University of Liverpool. NRF2^-/-^ A549 cells were generated using CRISPR/Cas9 technology as previously described [32].

### 2.3 CellTiter-Glo cell viability assay

Cells were seeded in 96-well plates at 5×10^4^ cells/well to achieve 70-80% confluency upon drug treatments. Following desired incubation periods, cells were lysed in 100 µL somatic cell ATP releasing reagent (Sigma, USA) before 1:1 dilution of cell lysate with 20 µL CellTiter-Glo Luminescent Cell Viability Assay (Promega, USA) as described by the manufacturer’s instructions. Luminescence was hereafter detected using a GloMax Multi-Detection System (Promega, USA). Bradford assays (Sigma, USA) were performed using cell lysates alongside a bovine serum albumin (BSA) standard curve for normalisation of ATP to protein content; absorbance was measured at 595 nm using a GloMax Multi-Detection System (Promega, UK). Cell viability was thus determined as ATP/mg protein as reported previously [25].

### 2.4 Quantitative real-time PCR

Total RNA was isolated from cell monolayers using the Monarch Total RNA Miniprep Kit (NEB, USA) according to the manufacturer’s instructions. cDNA was synthesised using LunaScript RT SuperMix Kit (NEB, USA) as per the instructions. Quantitative PCR (qPCR) analysis was performed on a ViiA 7 Real-Time PCR System (Applied BioSystems, USA) using Luna Universal qPCR Master Mix (NEB, USA). *GAPDH* was used as a normalisation control. Downstream data and statistical analyses were then reported as relative gene expression according to 2^-ΔCt^. Primer sequences are detailed in Table S1.

### 2.5 Western blotting

Cells were lysed in radioimmunoprecipitation (RIPA) lysis buffer (Millipore, USA) supplemented with 1% cOmplete protease and PhosStop phosphatase inhibitors (Roche, Switzerland) on ice for 15 min. Cell lysates were centrifuged at 13,500 rpm for 5 min at 4°C. Protein concentrations were determined using the Pierce BCA Protein Assay Kit (ThermoFisher, UK) according to the manufacturer’s instructions. Lysates were denatured using NuPAGE LDS Sample Buffer (Invitrogen, USA) supplemented with NuPAGE Sample Reducing Agent (Invitrogen, USA) for 5 min at 95°C. Proteins were separated by electrophoresis on NuPAGE Novex 4-12% Bis-Tris Protein Gels (Invitrogen, USA) and then electrophoretically transferred to Amersham Protran 0.45 µm nitrocellulose membranes (GE Healthcare, USA) at 300 mA for 90 min. After blocking with 5% non-fat milk for 1 h at room temperature, immunoblotting was performed at 4°C overnight using the following antibodies: rabbit anti-NRF2 (Proteintech, USA; 1:1000 dilution), rabbit anti-KEAP1 (Proteintech, USA; 1:10,000 dilution), goat anti-NQO1 (Everest, UK; 1:1000 dilution), rabbit anti-TXNRD1 (Abcam, UK; 1:5000 dilution), rabbit anti-GCLM (Abcam, UK; 1:10,000 dilution). Anti-α-Tubulin (Sigma, USA; 1:2000 dilution) was used as a loading control. Fluorescent secondary antibodies were purchased from LI-COR, USA: donkey anti-rabbit IRDye 800CW (1:25,000 dilution), donkey anti-mouse IRDye 800CW (1:25,000 dilution), donkey anti-rabbit IRDye 680RD (1:20,000 dilution), donkey anti-mouse IRDye 680RD (1:10,000 dilution), donkey anti-goat IRDye 800CW (1:25,000 dilution); all secondary antibody incubations were performed for 1 h at room temperature. Membranes were visualised using the Odyssey SA (LI-COR, USA).

### 2.6 LDH-based cytotoxicity assay

The CytoTox 96 Non-Radioactive Cytotoxicity Assay (Promega, UK) was utilised to measure LDH concentrations in both media and cell lysates according to the manufacturer’s instructions. Cells were seed in 96-well plates at 5×10^4^ cells/well and grown to 70-80% confluency following which cells were treated with CDDO ± DOX for the desired incubation periods. Lysis buffer was added to a set of untreated wells 45 min prior to the LDH assay to serve as positive controls for maximal cytotoxicity. Media was removed from each well and cells were lysed in lysis buffer. Both media and lysate samples were mixed 1:1 with CytoTox 96 reagent and incubated at room temperature for 30 min. Stop solution was added and absorbance read at 490 nm using a GloMax Multi-Detection System (Promega, UK). Released LDH (media) was quantified as a percentage of total LDH (media and lysate) to determine cytotoxicity.

### 2.7 Lentiviral transduction of AC16 cells

HEK293T cells (ATCC, USA) were transfected using Lipofectamine 2000 (ThermoFisher, UK) with a 3^rd^ generation lentiviral packaging system comprising the packaging plasmid (pRSV-Rev, Addgene, USA, #12253), envelope plasmid (pCMV-VSV-G, Addgene, USA, #8454) and the transfer plasmid containing the NRF2-mNeonGreen (mNG) sequence at a 2:2:1 ratio. Media containing lentiviral particles was collected after 48 h, pre-cleared by centrifugation and filtered through a 0.45 µm PES filter. For lentiviral transduction, AC16 cells were plated into 6-well plates with 7.5×10^5^ cells/well overnight. Subsequently, 5 mL undiluted lentivirus-rich supernatant was added for 72 h after which cells were imaged using an LSM780 microscope (Zeiss, Germany) to ensure 100% transduction efficiency. Cells were diluted into 96-well plates at a concentration of 50 cells/plate for monoclonal cell line generation to be used in downstream applications.

### 2.8 Live-cell imaging and fluorescence correlation spectroscopy (FCS)

AC16 cells were seeded into 4-compartment, glass-bottomed CellView 35 mm dishes (Greiner, Austria) at a density of 25,000 cells/well. Fluorescence imaging was performed using an LSM780 mounted onto an Axio Observer Z1 microscope with a 63x C-apochromat 1.4 NA oil-immersion objective lens (Zeiss, Germany); data were acquired via Zen 2012 software (Zeiss, Germany). The mNG fluorophore was excited using a 488 nm Argon laser with emission collected between 500-530 nm. Quantification of fluorescent counts was undertaken using previously described data collection and interpretation parameters [33]. Briefly, 5 x 2 s measurements were acquired using 0.2-0.4% laser power to optimise fluorescent signal with minimal photobleaching. Autocorrelation functions were derived automatically within the Zen 2012 software. Lambda scanning was performed on NRF2-mNG cells within the Zeiss software wherein data were acquired every 9 nm within the range 415-690 nm.

### 2.9 RNA interference (RNAi)

All RNAi reagents were purchased from Horizon Discovery, UK. AC16 cells (50,000 cells per well of a 6-well plate) were transfected with either SMARTpool non-targeting siRNA control, SMARTpool siRNA targeting *NFE2L2* or SMARTpool siRNA targeting *KEAP1,* diluted to desired amounts (10-50 ng) in siRNA buffer, using 4 µL of 20 µM DharmaFECT per well of a 6-well plate according to the manufacturer’s instructions. After 20 min at room temperature, fresh cell culture media was added. Total RNA was isolated 24-48 h later for qPCR analysis or protein extractions performed 48-72 h later for Western blotting to assess knockdown efficiency. SMARTpool siRNA sequences were as follows:

NFE2L2: UAAAGUGGCUGCUCAGAAU, GAGUUACAGUGUCUUAAUA, UGGAGUAAGUCGAGAAGUA, CACCUUAUAUCUCGAAGUU; KEAP1: GGACAAACCGCCUUAAUUC, CAGCAGAACUGUACCUGUU, GGGAGUACAUCUACAUGCA, CGAAUGAUCACAGCAAUGA; non-targeting: UGGUUUACAUGUCGACUAA, UGGUUUACAUGUUGUGUGA, UGGUUUACAUGUUUUCUGA, UGGUUUACAUGUUUUCCUA.

### 2.10 RNA-seq data analysis

RNA was extracted as described above. RNA-seq was performed by Novogene (Cambridge, UK) on a NovaSeq 6000 platform using a paired-end 150 bp (PE150) run configuration. Sequencing reads were aligned to the reference genome (GRCh38 primary genome assembly from Ensembl) using STAR [34]. Read counts for each gene (GRCh38 gene annotations from Ensembl) were generated using the featureCounts function of Subread [35]. Differential expression analysis was undertaken using R Bioconductor package DESeq2 [36] with filtering for effect size (log_2_ fold change >±0.58) and statistical significance (FDR <0.05). Volcano plots and heatmaps to display differential expression analysis were generated within Prism 10.4. Overrepresentation analysis (ORA) was performed using WEB-based Gene SeT AnaLysis Toolkit (WebGestalt) [37] in ORA mode using the gene ontology (GO) and Kyoto Encyclopedia of Genes and Genomes (KEGG) databases.

### 2.11 Statistical analysis

Data were analysed using Prism 10.0 software (GraphPad, USA). Normality tests were undertaken to determine the distribution of data. Statistical significance was p<0.05. One-way and two-way analysis of variance (ANOVA) were performed followed by Tukey’s or Dunnett’s post-hoc tests to evaluate statistical significance between multiple groups. Statistical differences between two groups were assessed through unpaired Student’s t-tests where significance was set at p<0.05.

## 3 Results

### 3.1 CDDO post-transcriptionally increases canonical NRF2 transcriptional target expression at both gene and protein levels

The maximal sub-toxic dose of CDDO was determined to be 100 nM, as established via cell viability assays (Fig. 1A). Thus, AC16 cardiomyocytes were treated with 100 nM CDDO for multiple timepoints ranging between 2-24 h. CDDO had no significant effect on *NFE2L2* mRNA expression at all timepoints (Fig. 1B). However, four well-established canonical NRF2 transcriptional targets (TXNRD1, NQO1, GCLM and FTH1) were significantly enriched 24 h following CDDO treatment (Fig. 1C-F). Interestingly, *TXNRD1* and *NQO1* mRNA levels were significantly upregulated after 4 h of CDDO treatment; *GCLM* and *FTH1* mRNA were significantly elevated above untreated control after 8 h (Fig. 1E-F). *KEAP1* expression was unaffected by CDDO treatment (Fig. 1G). Immunoblotting for NRF2, TXNRD1, GCLM and NQO1 revealed stabilisation of NRF2 protein significantly at 24 h and non-significantly at 72 h post-CDDO treatment (Fig. 1H-I). NRF2 protein was detected at ∼80 kDa and not the often reported 95-110 kDa due to the use of the NuPAGE Bis-Tris system with gradient gels and MOPS running buffer as has been highlighted previously with a suggested differential pH with respect to Tris-Glycine SDS-PAGE [38]. Downstream transcriptional targets of NRF2 were significantly enriched after 24 h (TXNRD1 and GCLM) and 72 h also (NQO1) (Fig. 1H-L). Hence, CDDO was capable of inducing NRF2 protein levels post-transcriptionally to cause increased activity of the transcription factor.

**Figure 1:**
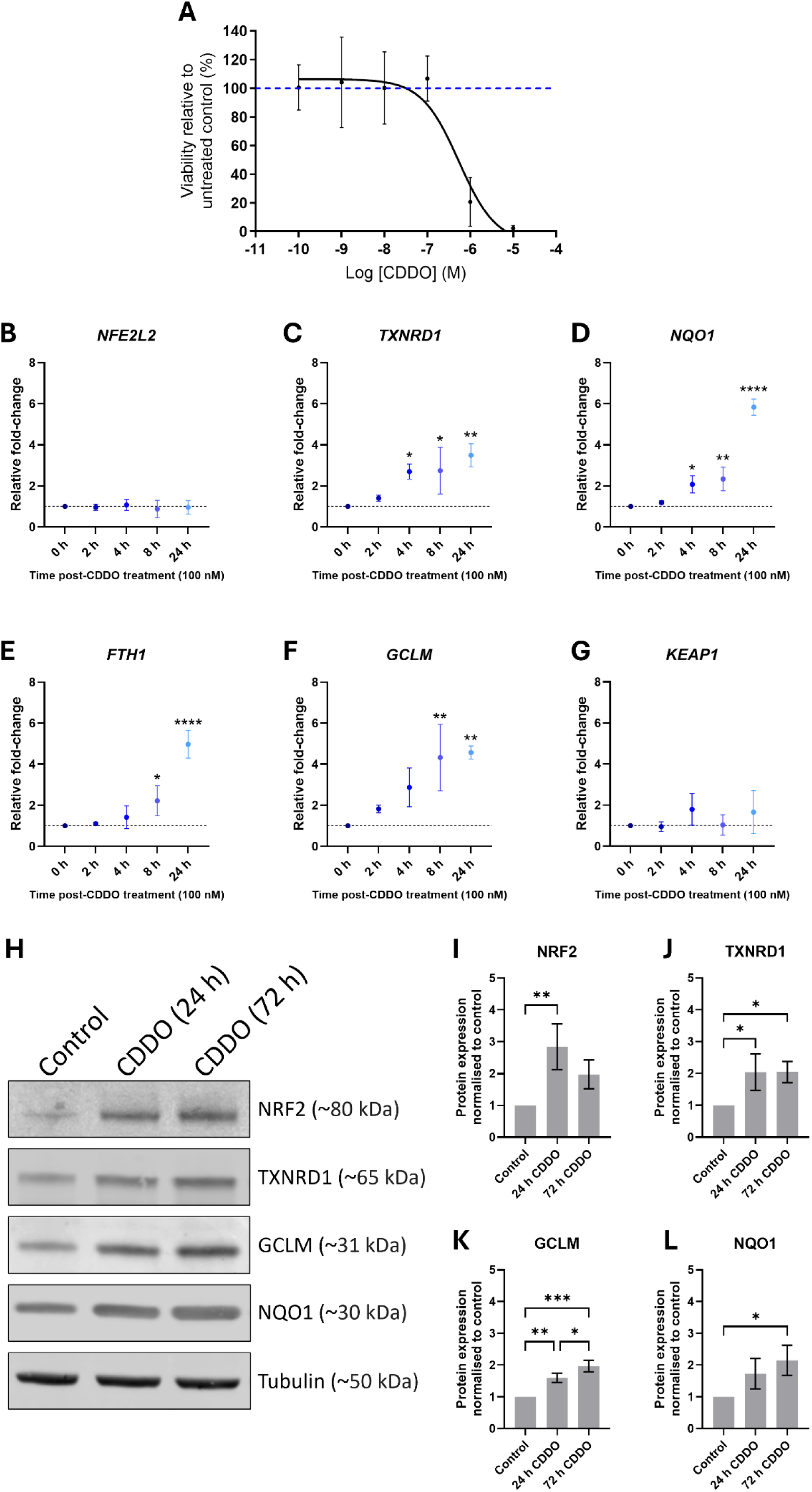
CDDO induces NRF2 transcriptional activity in AC16 cells. (**A**) Viability of AC16 cells in response to a CDDO concentration gradient with 72 h incubation, normalised to untreated control. Data are mean ± SD (n=3). Transcript levels of (**B**) *NFE2L2*, (**C**) *TXNRD1*, (**D**) *NQO1*, (**E**) *FTH1*, (**F**) *GCLM* and (**G**) *KEAP1* at numerous timepoints following 100 nM CDDO treatment. Data are mean ± SD (n=3). Statistical significance was determined using one-way ANOVAs with Tukey’s post-hoc analyses: *p<0.05, **p<0.01, ****p<0.0001 compared to untreated control. (**H**) Representative immunoblot and densitometric analyses of (**I**) NRF2, (**J**) TXNRD1, (**K**) GCLM and (**L**) NQO1 protein expression in WT AC16 cells treated with 100 nM CDDO (24 or 72 h). Data are mean ± SD (n=3) and normalised firstly to housekeeping protein α-tubulin then to untreated control cell lysates. Statistical significance was determined by one-way ANOVA with Tukey’s post-hoc analyses. *p<0.05, **p<0.01, ***p<0.001.

### 3.2 CDDO increases nuclear accumulation of NRF2 in AC16 cells

To quantify the subcellular localisation of NRF2 in response to CDDO, AC16 cells were transduced using a lentiviral approach to express NRF2-mNeonGreen (NRF2-mNG). A monoclonal cell line was produced on which fluorescence correlation spectroscopy (FCS) could be performed. Firstly, the fluorescent signal was confirmed to be mNG via lambda scanning where a peak intensity was measured at 521 nm, matching the known profile for mNG [39] (Fig. S1).FCS permitted the derivation of the NRF2-mNG molecule number within the confocal volume of the microscope. Within the cytoplasm, untreated NRF2-mNG AC16 cells expressed 5.77 ± 1.89 (mean ± SD) molecules/confocal volume which was significantly enriched (p<0.05) following 24 h incubation with 100 nM CDDO to 6.89 ± 2.12 molecules/confocal volume (Fig. 2A-B). Nuclear expression of NRF2-mNG under basal conditions (6.90 ± 2.73 molecules/confocal volume) was also significantly greater than untreated cytoplasmic expression (5.77 ± 1.89 molecules/confocal volume) and upregulated by a greater proportion with CDDO treatment, to 11.8 ± 3.88 molecules/confocal volume (p<0.0001) (Fig. 2B).

**Figure 2.**
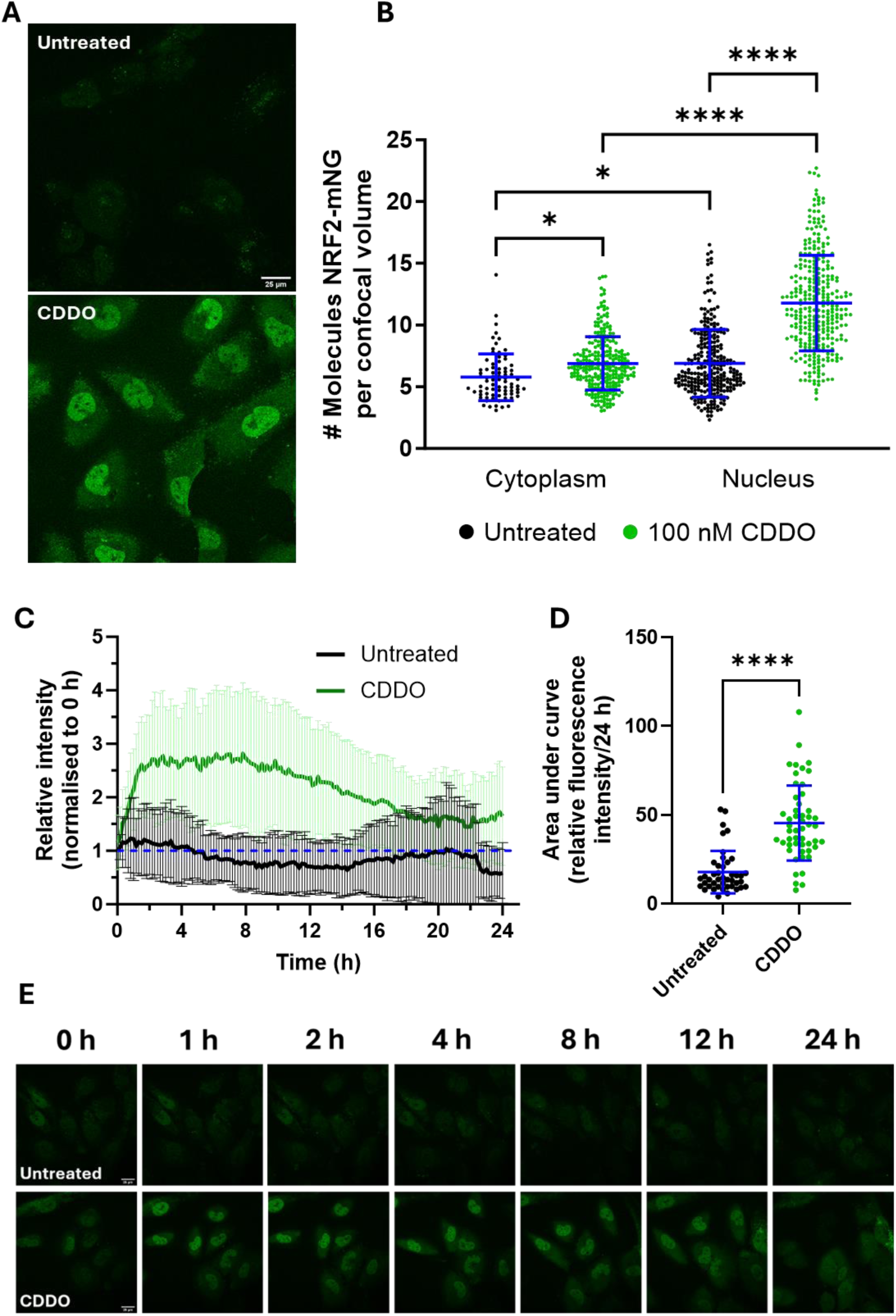
CDDO induces NRF2-mNeonGreen expression rapidly but sustained for 24 h. (**A**) Representative images of untreated and CDDO-treated (24 h, 100 nM) AC16 cells with stable expression of NRF2-mNG. (**B**) FCS-mediated quantification of nuclear or cytoplasmic NRF2-mNG molecule count per confocal volume. Data are mean ± SD (n≥67 individual cells from 3 independent experiments). *p<0.05, ****p<0.0001 following a one-way ANOVA with Tukey’s post-hoc analysis. (**C**) Timelapse tracking of nuclear NRF2-mNG signal in AC16 cells ± 100 nM CDDO with images acquired every 10 min for 24 h. Data are normalised to 0 h for each cell (n≥29 cells from 3 independent experiments) and reported as mean ± SD. (**D**) Area under the curve analysis of timelapse experiments. Data are mean ± SD. ****p<0.0001 following an unpaired, two-tailed Student’s t-test. (**E**) Representative images of timelapse experiment across 24 h. Scale bar indicates 25 µm.

Exposure to CDDO for 24 h was sufficient to increase NRF2 protein expression in AC16 cells. However, it has been well-established within the literature that NRF2 can be activated rapidly in response to accumulating ROS and has a short half-life. Thus, the NRF2-mNG-expressing AC16 cell line was utilised to measure dynamics of NRF2 following CDDO treatment in a time-lapse experiment across 24 h. Images from live cells were acquired every 10 min and nuclear NRF2-mNG fluorescence measured relative to 0 h, resulting in a fold-change in intensity (Fig. 2C-E). Following CDDO treatment, nuclear NRF2-mNG signal increased rapidly to 1.25-fold after 10 min, 1.6-fold after 30 min and 2-fold at 1 h post-treatment (Fig. 2C). Peak signal was achieved at ∼2 h post-CDDO at ∼3-fold increase from 0 h. This peak was sustained until 8 h post-treatment after which there was a slow and gradual decline towards baseline fluorescence measured at 0 h. After 24 h, however, fluorescence remained greater (∼1.7-fold) than baseline conditions, concurring with results presented above. Alternatively, non-CDDO treated AC16 cells showed relatively stable NRF2-mNG fluorescence within the nuclei. The area under the curve for nuclear fluorescence within each cell was calculated. CDDO-treated cells showed significantly greater nuclear NRF2-mNG signal with respect to untreated cells (p<0.0001, two-tailed, unpaired Student’s t-test) (Fig. 2D).

### 3.3 Doxorubicin induces dose-dependent cytotoxicity in AC16 cells

AC16 cells were incubated with variable DOX concentrations (seven concentrations within 100 pM – 100 μM) over a 48 h period (Fig. 3A). Cell viability was reduced in a dose-dependent manner following DOX treatment with mean IC_50_ measured as 117 nM. Cells were pre-treated with 100 nM CDDO for 24 h prior to DOX exposure to permit NRF2-mediated upregulation of downstream targets, as described earlier. When subsequently incubated with a DOX concentration gradient, the IC_50_ of CDDO-pretreated cells increased to 245 nM (Fig. 3A). Cell viability in response to 100 nM DOX was 44.0% (± 3.23), a significant reduction from untreated cells (p<0.0001) yet, with pre-exposure to CDDO for 24 h prior to DOX, this was significantly increased to 62.6% (± 6.73) (p<0.05) (Fig. 3B).

**Figure 3:**
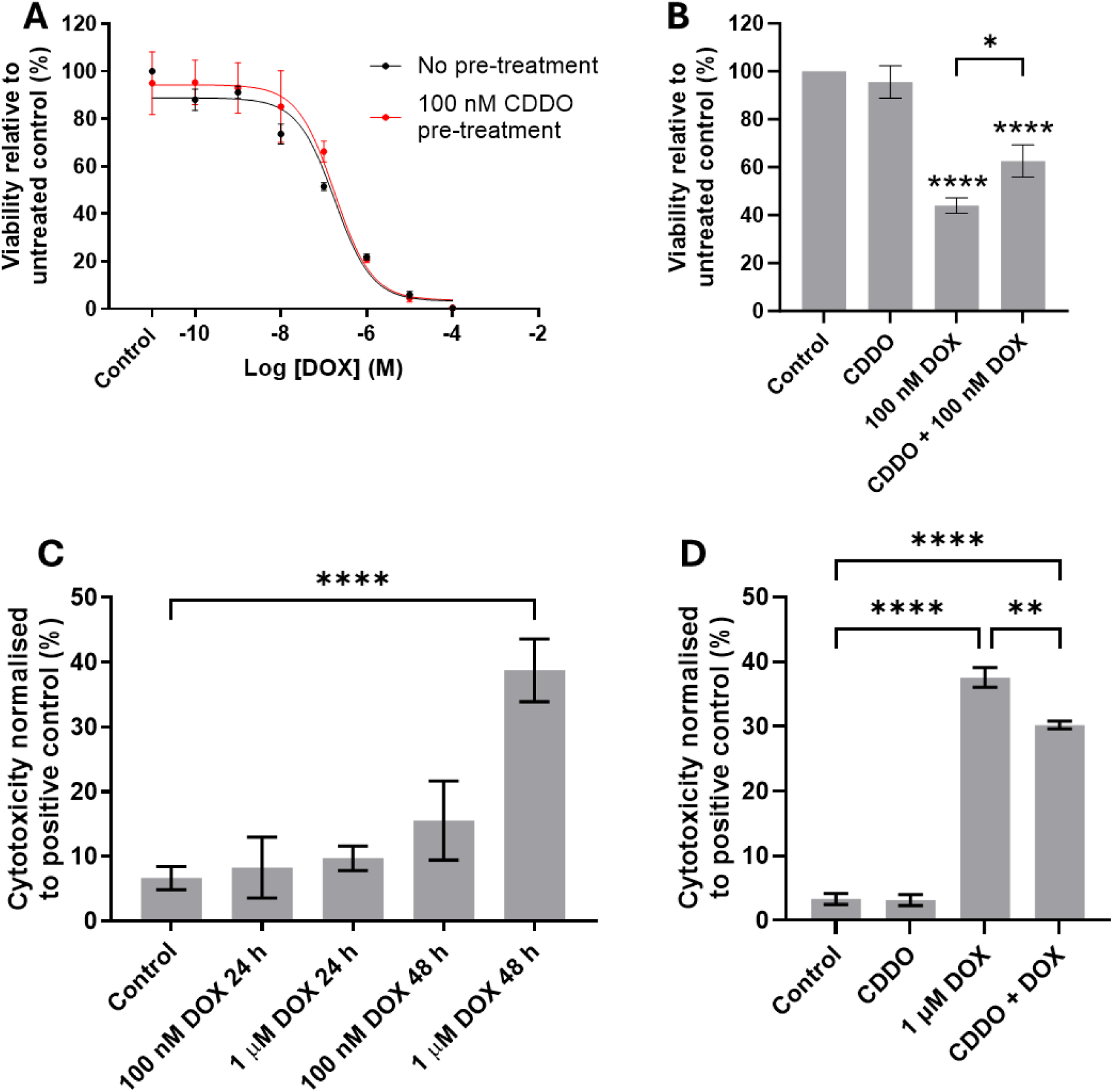
CDDO can protect AC16 cells from DOX-induced toxicity. (**A**) AC16 cell viability in response to 48 h incubation with a DOX concentration gradient ± 24 h pre-treatment with 100 nM CDDO. Data are normalised to untreated controls and presented as mean ± SEM (n=3). IC_50_ with no pre-treatment was 117 nM (95% CI: 67.6 – 203 nM); IC_50_ with 100 nM CDDO pre-treatment was 245 nM (95% CI: 110 – 558 nM); comparison of nonlinear fits suggested a non-significant effect (p=0.13) following a sum-of-squares F test. (**B**) Cell viability in response to 100 nM DOX (48 h) ± 24 h pre-treatment with 100 nM CDDO. Data are mean ± SEM (n=6). (**C**) Cytotoxicity of AC16 cells in response to variable DOX doses and incubation times following an LDH assay. Data are normalised to a 100% cytotoxic control and presented as mean ± SEM (n≥2). (**D**) AC16 cell cytotoxicity in response to 1 µM DOX (48 h) ± 24 h pre-incubation with 100 nM CDDO. Statistical significance was determined following one-way ANOVAs with Tukey’s post-hoc analyses; *p<0.05, **p<0.01, ****p<0.0001.

Cytotoxicity of DOX in AC16 cells was also evaluated via LDH assays with released LDH calculated as a proportion of total LDH (from media and cell lysates to account for any LDH decay within the media) and normalised to a positive control (100% cytotoxicity). Firstly, the dose and timepoint for DOX incubation was optimised wherein only 1 µM DOX for 48 h induced a significant increase (p<0.0001) in toxicity within the assessed treatment range (Fig. 3C) from 6.65% (± 1.80) in control cells to 38.8% (± 2.81) in 1 µM 48 h DOX. Pre-treatment of AC16 cells with 100 nM CDDO for 24 h prior to 1 µM DOX (48 h) was able to significantly reduce the cytotoxicity of the chemotherapeutic (Fig. 3D).

### 3.4 Genetic overexpression of NRF2-mNG in AC16 cells promotes cardioprotection against DOX

A consequence of using the lentiviral expression approach for transduction of AC16 cells is the opportunity for multiple integrations of the plasmid into the host genome resulting in overexpression of the fluorescently tagged NRF2 above endogenous NRF2 levels. To ascertain if this was successfully achieved, transcript levels of NRF2 and target genes were assessed via qPCR (Fig. 4A). The NRF2-mNG expressing cell line exhibited significantly greater *NFE2L2* levels and canonical NRF2 target genes *NQO1*, *FTH1* and *GCLM* than the wild-type (WT) AC16 cell line; *KEAP1* and *TXNRD1* transcript levels were not significantly different (Fig. 4A). Interestingly, despite the increased basal expression, *FTH1* and *GCLM* expression was significantly induced by CDDO relative to untreated cells (Fig. 4A). As previously established, *NFE2L2* transcript levels were unaffected by CDDO treatment (Fig. 4A). Moreover, *NQO1* was not induced by CDDO treatment in NRF2-mNG expressing AC16 cells, in contrast to WT AC16 (Fig. 1D). NRF2 protein levels were subsequently measured via Western blotting. Endogenous NRF2 was observed at ∼80 kDa in both cell lines which was significantly increased following 24 h CDDO and 2 h MG132 treatment (a proteasomal inhibitor that can be utilised for NRF2 stabilisation [32]) in the WT AC16 cell (p<0.05) but not in the NRF2-mNG AC16 cells (Fig. 4B-C). The molecular weight of the mNG tag is ∼27 kDa [39] therefore NRF2-mNG was predicted to appear at ∼107 kDa. Strong bands were observed in the NRF2-mNG AC16 cell line at this molecular weight which were CDDO and MG132-inducible (Fig. 4B, D).

**Figure 4:**
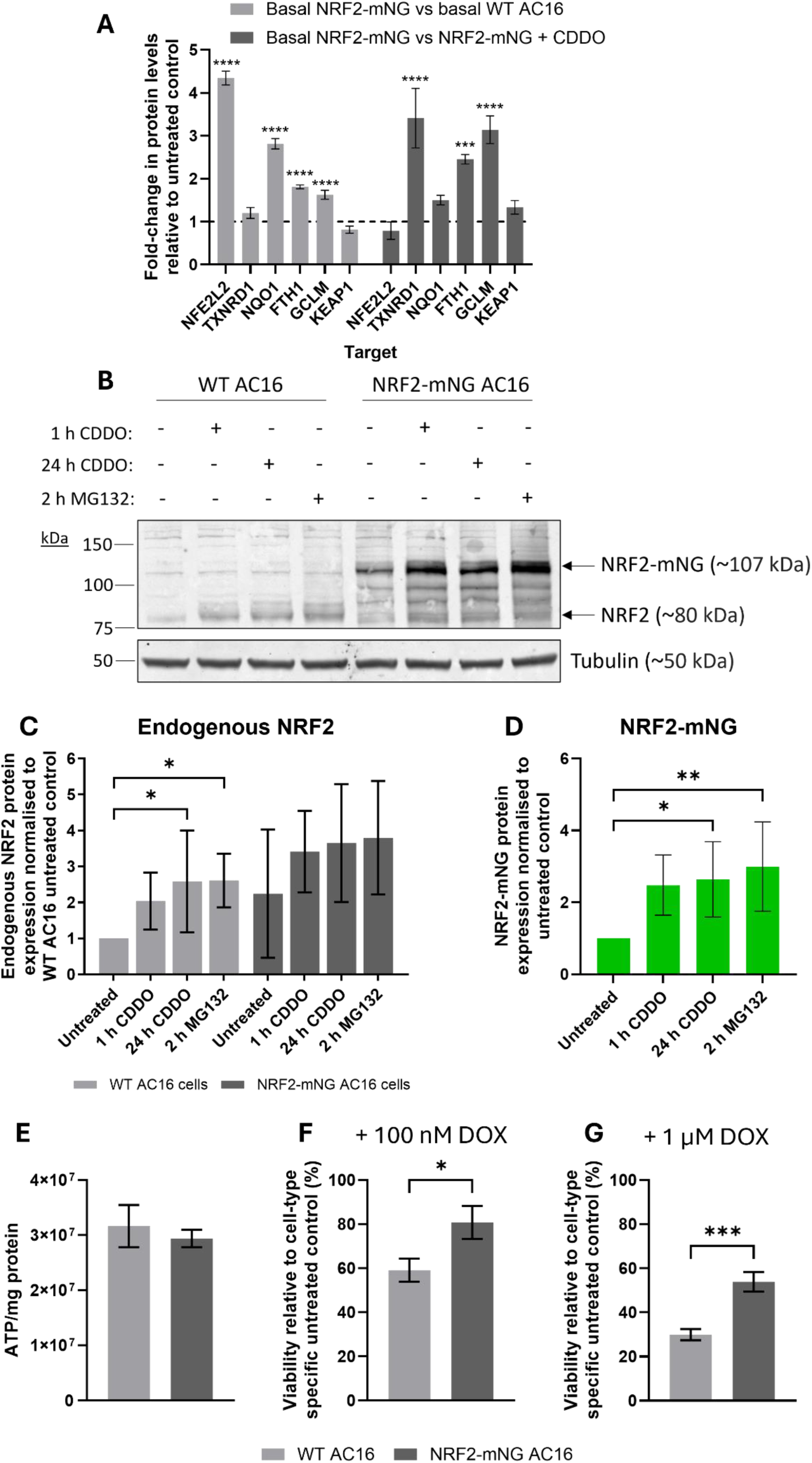
Overexpression of NRF2-mNG confers protection of AC16 cells to doxorubicin cytotoxicity. (**A**) Transcript expression of *NFE2L2*, canonical NRF2-target genes (*TXNRD1*, *NQO1*, *FTH1* and *GCLM*) and *KEAP1* in AC16 cells expressing NRF2-mNG, relative to WT AC16 cells under basal and 24 h CDDO-stimulated conditions. Data are expressed as mean ± SD (n=3 independent experiments). Statistical difference was evaluated using one-way ANOVA with Dunnett’s correction; ***p<0.001, **** p<0.0001 compared to relative untreated or CDDO-treated WT AC16 controls. (**B**) Representative immunoblot and (**C**) densitometric analysis for endogenous NRF2 (∼80 kDa) in WT and NRF2-mNG-expressing AC16 cells treated with 100 nM CDDO (1 or 24 h), 10 µM MG132 (2 h) or left untreated prior to lysis. Data are mean ± SD (n=5 independent repeats) and normalised to WT AC16 untreated control. (**D**) Densitometric analysis of NRF2-mNG in the NRF2-mNG-expressing AC16 cell line following treatment with 100 nM CDDO (1 or 24 h), 10 µM MG132 (2 h) or left untreated prior to lysis. Data are mean ± SD (n=5 independent repeats) and normalised to untreated NRF2-mNG levels. Statistical differences were determined using a one-way ANOVA with Dunnett’s correction. (**E**) Basal ATP content of cell lysates from WT and NRF2-mNG AC16 cells relative to protein content. Data are mean ± SEM (n≥5). (**F-G**) Cell viability of WT and NRF2-mNG AC16 cells upon 48 h treatment with 100 nM or 1 µM DOX. Data are mean ± SEM (n=8) and normalised to cell-type specific untreated controls. Statistical significance was evaluated using unpaired, two-tailed Student’s t-tests; *p<0.05, ***p<0.001.

The functional consequence of NRF2-mNG genetic overexpression in AC16 cells with respect to DOX sensitivity was assessed. It was also important to evaluate whether the NRF2-mNG molecules could impact the cytoprotective nature of NRF2. Firstly, basal ATP production was measured with no significant difference between cell types (Fig. 4E). Both WT and NRF2-mNG AC16 cells were then incubated with 100 nM and 1 µM DOX for 48 h. Cell viability of the NRF2-mNG expressing cells was significantly increased relative to WT cells when incubated with both DOX concentrations (Fig. 4F-G).

### 3.5 Transient knockdown of KEAP1 induces NRF2 activity in AC16 cells

To further evaluate the contribution of NRF2 to protection against DOX cytotoxicity, genetic upregulation and perturbation of NRF2 activity was explored using *NFE2L2* and *KEAP1* siRNA, respectively. To optimise siRNA transfection conditions, AC16 cells were transfected with 10, 25 or 50 ng of siRNA targeting *NFE2L2* and *KEAP1* for 24 and 48 h, followed by quantification of transcript levels using qPCR (Fig. S2). All three siRNA concentrations effectively reduced target gene expression at both time points, with no significant differences observed between the conditions (Fig. S2A–B). Pooled data across concentrations and timepoints thus highlighted that control, non-targeting siRNA had no effect on *NFE2L2* or *KEAP1* levels (Fig. S2C-D); *NFE2L2* mRNA was knocked down by ∼92% (p<0.01) (Fig. S2C) and *KEAP1* mRNA was knocked down by ∼89% (p=0.07) (Fig. S2D).

After this, efficacy of the siRNAs for protein knockdown was optimised via Western blotting. Cells were transfected with 10 or 25 ng siRNA for 48 h followed by a further 24 h incubation with CDDO (Fig. 5). As before, CDDO significantly increased NRF2 protein expression in non-transfected AC16 cells (p<0.05) (Fig. 5B). Moreover, GCLM and NQO1 expression was significantly increased (p<0.05 and p<0.0001, respectively) upon 24 h CDDO treatment (Fig. 5D-E). Control, non-targeting siRNA transfection had no significant impact on the expression of NRF2, KEAP1, GCLM or NQO1 under basal conditions and, whilst the stimulative impact of CDDO on NRF2 and GCLM levels were non-significant, NQO1 significant upregulation was maintained (p<0.0001). *NFE2L2* siRNA inhibited NRF2 protein levels to 35.0 ± 0.06% of non-transfected control cells (Fig. 5B). However, these effects were more apparent when comparing CDDO-treated lysates where NRF2 was significantly knocked down by *NFE2L2* siRNA (p<0.001) compared to control (Fig. 5B). This effect was also measured for GCLM and NQO1 with significant inhibition under a CDDO treated background (p<0.05 and p<0.0001, respectively; Fig. 5D-E). *KEAP1* siRNA significantly downregulated KEAP1 protein expression (p<0.0001) to 42.3 ± 0.18% that of non-transfected cells (Fig. 5C) but interestingly with no significant effect on NRF2 protein under basal conditions (Fig. 5B). CDDO was also able to significantly increase NRF2 protein even alongside *KEAP1* siRNA (p<0.05), comparably to control cells (Fig. 5B). Whilst GCLM protein was not significantly affected by KEAP1 suppression, NQO1 was significantly upregulated to 3.23 ± 1.15-fold greater than non-transfected cells in untreated conditions (p<0.01). MG132 (10 µM, 2 h) was also utilised for NRF2 stabilisation (Fig. S3); these results corroborated these data wherein *NFE2L2* siRNA reduced NRF2 expression and KEAP1 protein was diminished upon *KEAP1* siRNA transfection to a significant degree (p<0.01, p<0.001, respectively) after 10 ng siRNA transfection notably. It was thus determined that both *NFE2L2* and *KEAP1* siRNA were having a direct impact on the expression of the respective target proteins. Subsequently, the functional consequences of these protein manipulations were assessed with regards to DOX sensitivity.

**Figure 5:**
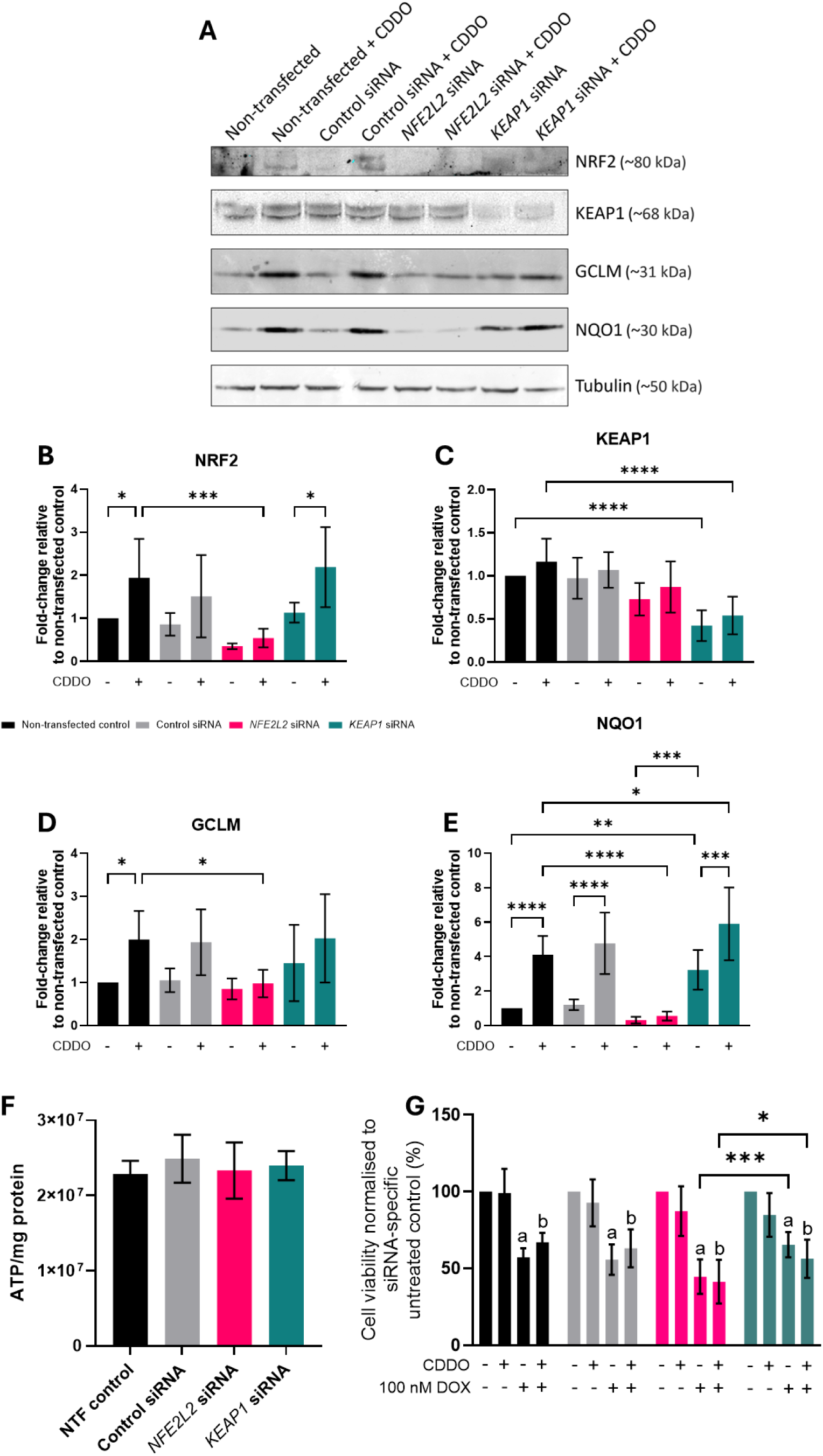
Genetic knockdown of NRF2 increases AC16 sensitivity to DOX whereas genetic upregulation of NRF2 provides cytoprotection. (**A**) Representative immunoblots with AC16 cells 72 h post-transfection with control, non-targeting siRNA, *NFE2L2* siRNA or *KEAP1* siRNA ± 24 h CDDO treatment (100 nM). (**B-E**) Quantification of NRF2, KEAP1, GCLM and NQO1 protein expression. Data are mean ± SD (n=6) and normalised to untreated, non-transfected AC16 cells. Statistical significance was evaluated using one-way ANOVAs with Tukey’s post-hoc analyses. (**F**) Basal ATP production in AC16 cells following siRNA transfections; NTF, non-transfected. (**G**) Cell viability in siRNA-transfected AC16 cells in response to 100 nM DOX (48 h) ± 24 h pre-treatment with 100 nM CDDO. Data are normalised to transfection-specific untreated AC16 cells and presented as mean ± SEM (n=8). Statistical significance was determined using a two-way ANOVA with Tukey’s post-hoc analysis; a and b indicate p<0.0001 with respect to untreated (a) or CDDO-treated cells (b), respectively. *p<0.05, ***p<0.001 between specific groups.

### 3.6 NRF2 contributes to CDDO-mediated AC16 protection

Firstly, it was confirmed that ATP production of AC16 cells in response to the various siRNA transfections was unchanged (Fig. 5F). DOX treatment (100 nM, 48 h) significantly reduced cell viability under all siRNA backgrounds (Fig. 5G). Whilst no significant difference was detected between non-transfected and *NFE2L2* or *KEAP1* downregulation in response to DOX, it was notable that direct comparison of these two groups revealed a significant (p<0.001) difference directly between them whereby *KEAP1* siRNA-transfected cells were less sensitive to DOX than *NFE2L2* siRNA (Fig. 5G). The impact of NRF2 on the protection afforded to AC16 cells via CDDO treatment as shown previously was thus analysed via siRNA treatments. Whilst cytoprotection with CDDO + DOX relative to DOX-treated was measured in AC16 cells (Fig. 3B), any cytoprotection was ablated under *NFE2L2* and *KEAP1* siRNA backgrounds (Fig. 5G). This suggests that CDDO may require functional NRF2 to provide a protective phenotype against DOX cytotoxicity in AC16 cells.

### 3.7 CDDO did not promote DOX resistance in cancer cells

The A549 lung adenocarcinoma cell line has a somatic mutation within *KEAP1* resulting in constitutive activation of NRF2 and promoting chemotherapeutic resistance [40,41]. Thus, an NRF2^-/-^ A549 cell line was produced using CRISPR/Cas9 technology to measure the impact on DOX sensitivity. Firstly, NRF2 and target protein expression were analysed by immunoblotting (Fig. 6A). CDDO treatments for 2 and 24 h alongside proteasomal inhibition through MG132 were used to stabilise NRF2 prior to cell lysis. Firstly, NRF2 was not enriched upon CDDO treatment in WT A549 cells and was not significantly upregulated with MG132 despite a tendency towards an increased expression (Fig. 6B). The NRF2^-/-^ genotype was validated with qualitative ablation of NRF2 protein bands using immunoblotting and quantitatively significant reduction of NRF2 (p<0.01) with respect to all WT A549 untreated and CDDO or MG132-treated conditions (Fig. 6B). TXNRD1, GCLM and NQO1 expression were also unaffected by treatment in both cell lines although were significantly diminished in the NRF2^-/-^ A549 when contrasted with WT cells (Fig. 6C-E).

**Figure 6:**
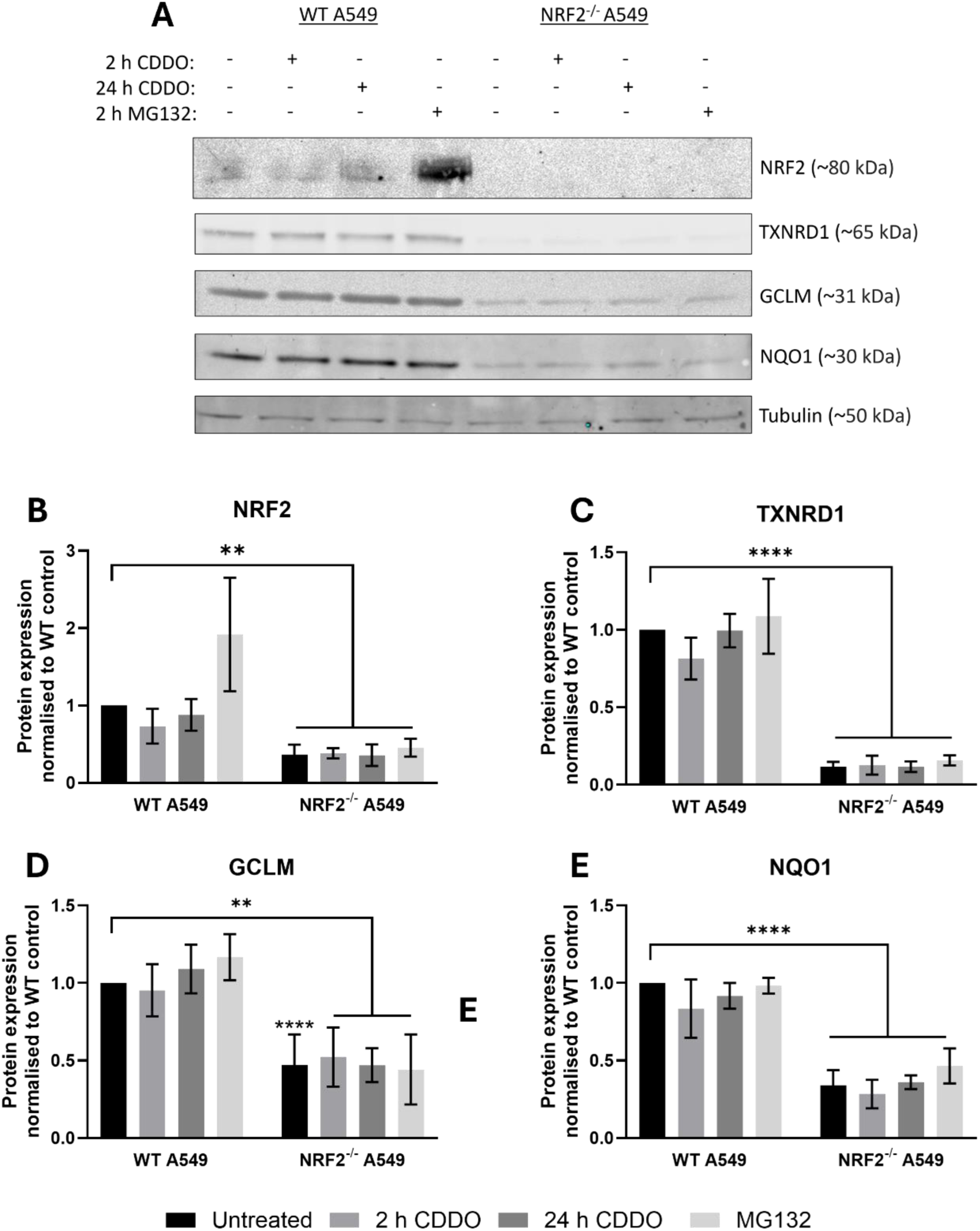
Generation and quantification of NRF2^-/-^ A549 lung adenocarcinoma cells. (**A**) Representative immunoblot of A549 cells with CRISPR/Cas9-mediated NRF2 double-knockout or WT genotypes ± 100 nM CDDO (2 or 24 h) or 10 µM MG132 (2 h). (**B-E**) Quantification of NRF2, TXNRD1, GCLM and NQO1 protein expression. Data are mean ± SD (n=3) and normalised to untreated, WT control A549 cells. Statistical significance was evaluated using one-way ANOVAs with Tukey’s post hoc analyses. **p<0.01, ****p<0.0001.

Basal ATP/protein was not significantly impacted by NRF2 ablation in A549 cells (Fig. 7A). In WT cells, 100 nM DOX did not significantly diminish cell viability although was reduced to 77.6% (± 4.78) whereas 1 μM DOX resulted in significant reduction (p<0.0001) to 39.2% (± 9.79) (Fig. 7B). CDDO pre-treatment for 24 h (which showed significant cytoprotection in AC16 cells; Fig. 2B) exacerbated the cytotoxicity of 100 nM DOX; cell viability after CDDO + 100 nM DOX was significantly reduced with respect to untreated cells (p<0.05) whereas DOX treatment by itself did not significantly change viability (Fig. 7B). Viability of NRF2^-/-^ A549 cells was significantly reduced to 47.6% (± 9.85) and 23.5% (± 14.1) following 48 h incubation with 100 nM (p<0.001) and 1 μM DOX (p<0.0001), respectively (Fig. 7C). CDDO pre-treatment did not increase DOX toxicity but had no beneficial effect on cell viability in these cells (Fig. 7C). Direct comparison between A549 genotypes following 100 nM DOX treatment revealed significantly reduced cell viability in the NRF2^-/-^ cell line with respect to WT (p<0.01) (Fig. S4A); no significant effect was measured in response to 1 μM DOX (Fig. S4B). LDH assays also highlighted that 48 h treatment with either 100 nM or 1 μM DOX had no impact on WT A549 cells (Fig. 7D). However, DOX induced a dose-dependent effect in NRF2^-/-^ A549 cells (p<0.0001) showing a significant increase in cytotoxicity after 1 µM treatment compared to untreated cells (Fig. 7E). CDDO also showed no cytoprotective properties via the LDH assay regardless of A549 genotype (Fig. 7D-E).

**Figure 7:**
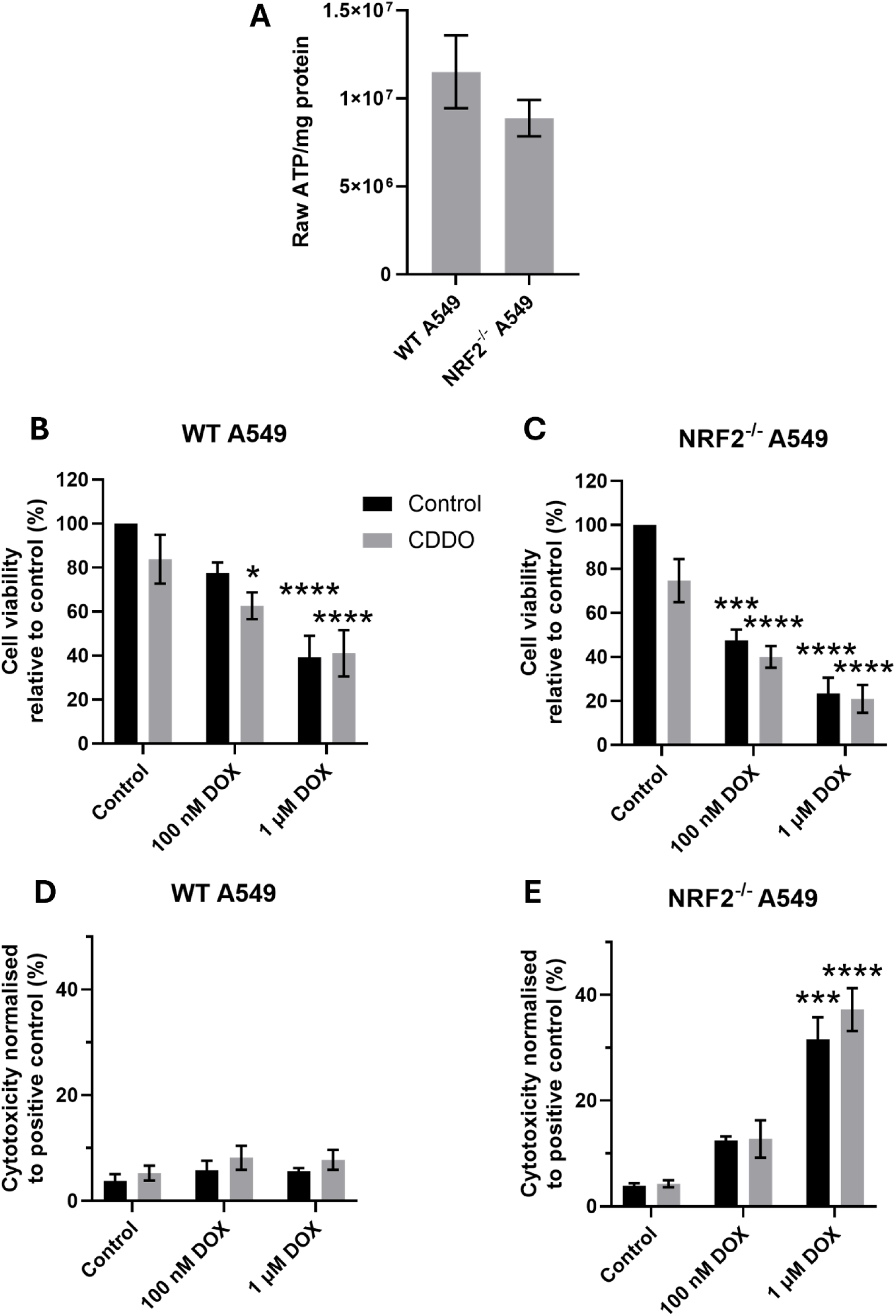
NRF2^-/-^ A549 cells are more sensitive to DOX cytotoxicity than WT. (**A**) Basal ATP production in WT and NRF2^-/-^ A549 cells. (**B-C**) Cell viability and (**D-E**) cytotoxicity in WT and NRF2^-/-^ A549 cells, respectively, in response to 48 h DOX treatment (100 nM and 1 µM) ± 24 h pre-incubation with 100 nM CDDO. Data are normalised to genotype-specific untreated controls (B-C) or a 100% cytotoxic control (D-E) and presented as mean ± SEM (n≥3). Statistical significance was determined following a two-way ANOVA with Tukey’s post-hoc analyses. *p<0.05, ***p<0.001, ****p<0.0001 relative to untreated controls.

Two pancreatic cancer cell lines were also utilised to assess the impact of NRF2 induction for potential DOX resistance; SUIT-2 and MIA PaCa2. These were selected based upon previous evidence that NRF2 overexpression in pancreatic cancer and these cell lines specifically can promote chemotherapeutic resistance, perhaps similarly to A549 [42].Firstly, the expression and activity of NRF2 was assessed (Fig. 8A); basal expression of NRF2 and canonical NRF2 transcriptional targets TXNRD1, GCLM and NQO1 were compared. NRF2 and GCLM were not significantly different between the two cell lines (Fig. S5). However, MIA PaCa2 cells had significantly downregulated TXNRD1 expression (p<0.05) yet significantly greater NQO1 levels (p<0.0001) at 3.89-fold (± 0.26) with respect to SUIT-2 cells (Fig. S5). Lysates were collected from both cell lines after CDDO (2 and 24 h) and MG132 (2 h) treatment for the stabilisation of NRF2. NRF2 levels were not significantly affected by these treatments in SUIT-2 cells (Fig. 8B). However, TXNRD1 and GCLM protein expression was significantly enhanced after 24 h CDDO relative to untreated cells (p<0.01 and p<0.001, respectively); NQO1 levels were unaffected (Fig. 8C-E). MIA PaCa2 cells treated with CDDO for both 2 and 24 h or MG132 for 2 h showed significantly enriched NRF2 (p<0.01 for 2 h CDDO and p<0.001 for 24 h CDDO and 2 h MG132) (Fig. 8F). However, the expression of all other proteins remained unchanged (Fig. 8G-I).

**Figure 8:**
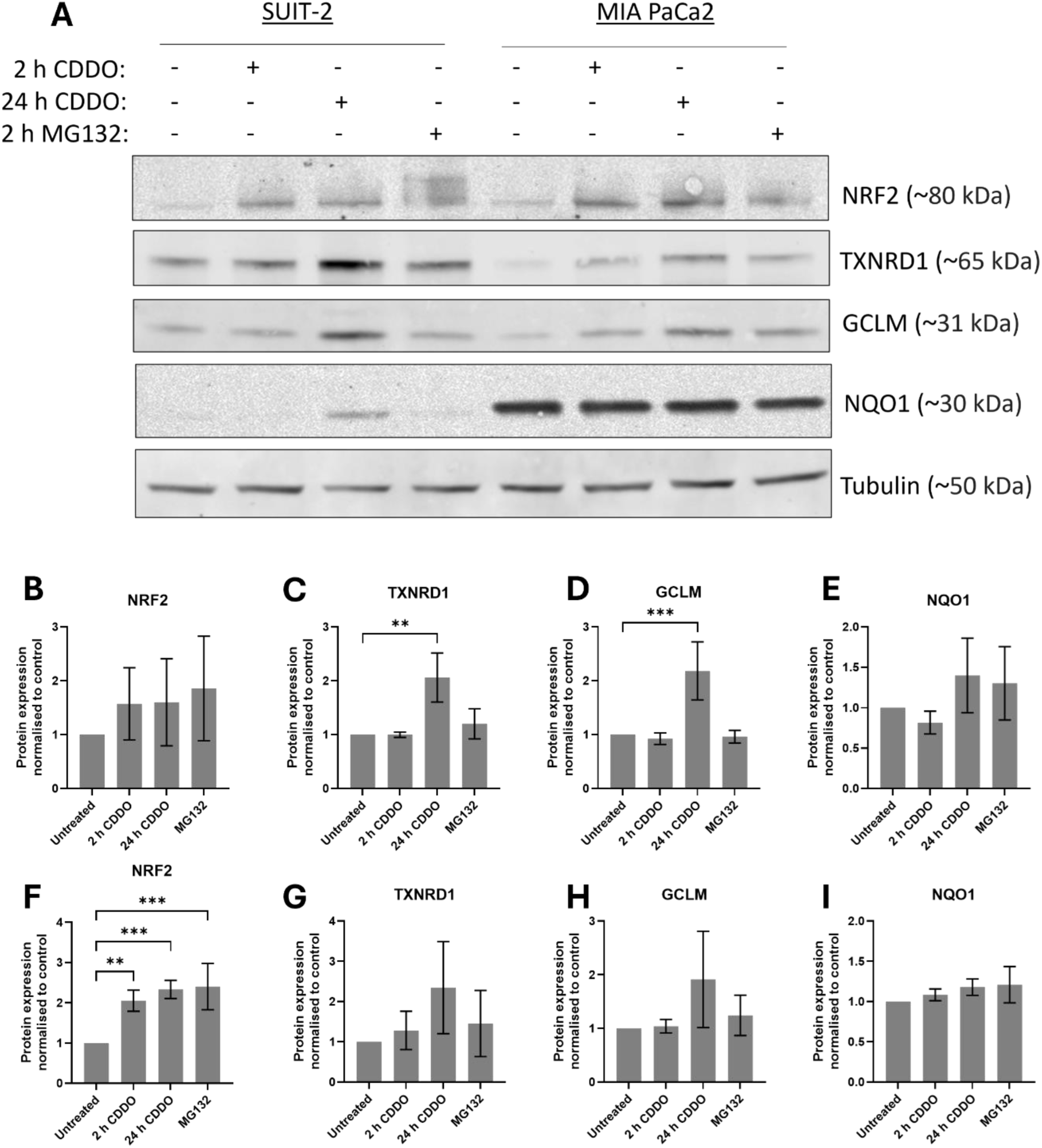
Assessment of NRF2 and NRF2-target protein expression in SUIT-2 and MIA PaCa2 pancreatic cancer cell lines. (**A**) Representative immunoblot of SUIT-2 and MIA PaCa2 cells ± 100 nM CDDO (2 or 24 h) or 10 µM MG132 (2 h). (**B-E**) Quantification of NRF2, TXNRD1, GCLM and NQO1 protein expression in SUIT-2 cells and (**F-I**) MIA PaCa2 cells, respectively. Data are normalised to cell type-specific untreated controls and reported as mean ± SD. Statistical significance was determined via one-way ANOVAs with Tukey’s post-hoc analyses. **p<0.01, ***p<0.001.

The functional effect of CDDO and DOX within these two cell types was subsequently assessed in cell viability and cytotoxicity assays. Basal ATP/protein was comparable between the cell lines (Fig. 9A). Moreover, DOX dose-dependently and significantly reduced cell viability in both cell lines (Fig. 9B-C). Direct comparison between SUIT-2 and MIA PaCa2 cell viability following 48 h treatment with 100 nM and 1 µM DOX showed analogous results (Fig. S4C-D). As in A549 cells, CDDO had no protective impact when pre-treated for 24 h prior to DOX delivery; CDDO may have promoted the impact of DOX in reducing cell viability in MIA PaCa2 cells exposed to 100 nM DOX (Fig. 9C). Interestingly, neither DOX nor CDDO significantly affected cytotoxicity in either pancreatic cancer cell line following LDH assays (Fig. 9D-E).

**Figure 9:**
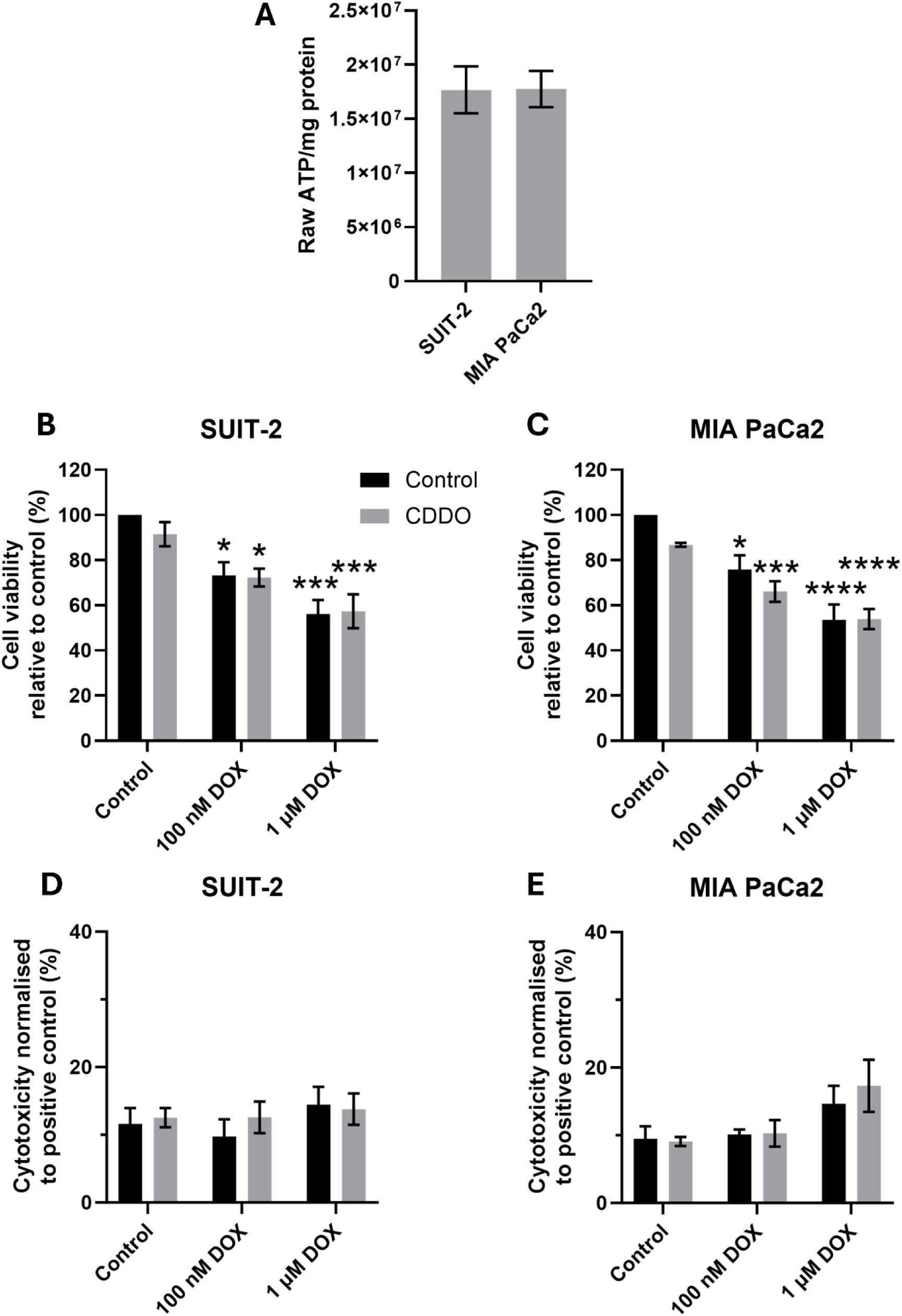
CDDO does not promote SUIT-2 or MIA PaCa2 survival in the presence of DOX. (**A**) Basal ATP production in SUIT-2 and MIA PaCa2 cells. (**B-C**) Cell viability and (**D-E**) cytotoxicity in SUIT-2 and MIA PaCa2 cells, respectively, in response to 48 h DOX treatment (100 nM and 1 µM) ± 24 h pre-incubation with 100 nM CDDO. Data are normalised to genotype-specific untreated controls (B-C) or a 100% cytotoxic control (D-E) and presented as mean ± SEM (n≥3). Statistical significance was determined following a two-way ANOVA with Tukey’s post-hoc analyses. *p<0.05, ***p<0.001, ****p<0.0001 relative to untreated controls.

### 3.8 CDDO specifically upregulates NRF2-related genes to drive cytoprotective pathways in the AC16 transcriptome

Considering the suggestion that CDDO alleviates AC16 cytotoxicity from DOX in an NRF2-dependent manner, but not that from cancer cells, the transcriptomic profiling of AC16 cells after CDDO treatment was performed. Samples were analysed after 24 h of CDDO incubation to ascertain transcriptional effects after the pre-treatment period prior to DOX treatment in results presented thus far and after 72 h CDDO treatment to mimic the full incubation window. Moreover, this provided insight into acute and more sustained impacts of CDDO treatment. A total of 76 significantly differentially expressed genes (DEGs) were measured 24 h after CDDO treatment of which 49 were upregulated and 27 downregulated with respect to control, vehicle (DMSO)-treated samples (Fig. 10A). The number of DEGs was enhanced with longer CDDO exposure with 277 DEGs in total (147 upregulated and 130 downregulated) (Fig. 10B). Comparison of these DEG profiles between the two time points revealed strong overlap wherein 44/49 upregulated DEGs after 24 h continued to be significantly enriched at 72 h (Fig. 10C). Similarly, 13/27 downregulated DEGs after 24 h CDDO treatment relative to control samples were also downregulated at 72 h (Fig. 10C). Notably, 103 and 117 DEGs were uniquely upregulated and downregulated, respectively, by 72 h treatment that were not significantly changed by 24 h CDDO treatment (Fig. 10C). A manually curated list of 53 canonical and consensus NRF2 target genes was produced based upon literature evidence [43–45] (Table S2) which encompassed genes encoding proteins involve with multiple biological roles including biotransformation, detoxification, anti-oxidation, metabolism, bioenergetics and inflammation. After 24 h CDDO treatment, none were downregulated and 17 were significantly enriched DEGs; by 72 h of CDDO incubation, 26/53 were significantly enriched (Fig. 10D). At both timepoints, *HMOX1*, *HKDC1*, *SLC7A11* and *NQO1* were the top 4 most upregulated genes. Transcriptional expression of *NFE2L2* itself was unaffected by CDDO treatment regardless of timepoint, demonstrating consistency with qPCR data (Fig. 1B, Fig. 2A).

**Figure 10:**
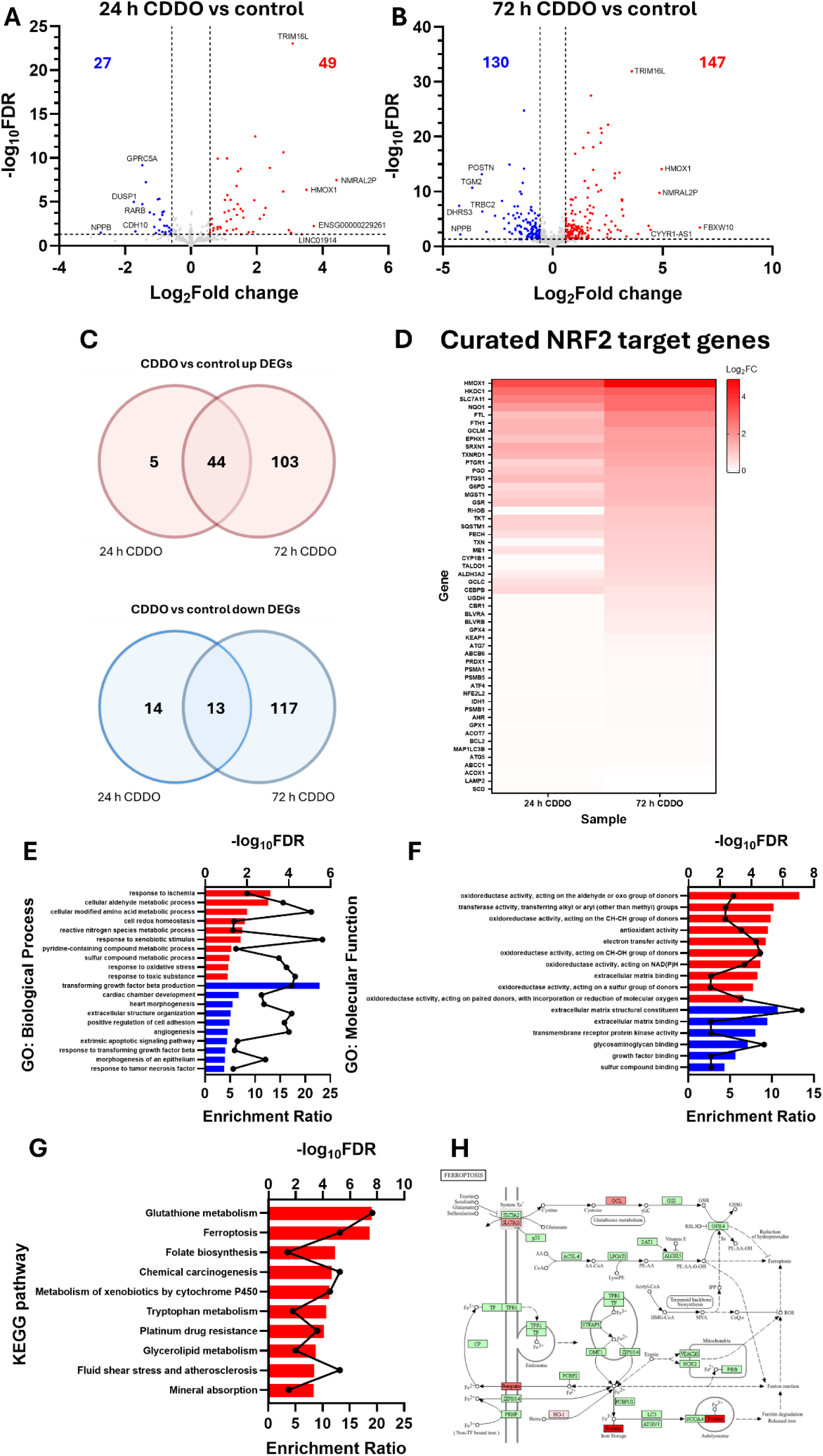
CDDO specifically activates NRF2-related genes pathways after 24 and 72. **h.** (**A-B**) Differential expression analysis volcano plots for 24 h and 72 h CDDO incubation (100 nM) compared to vehicle-treated control (n=3 biological repeats for all conditions). Blue and red points indicate downregulated and upregulated DEGs, respectively. DEGs were defined by log_2_FC≥0.58 or ≤-0.58 and FDR≤0.05 (**C**) Overlap of 24 h and 72 h upregulated (red) or downregulated (blue) DEGs relative to vehicle-treated control in AC16 cells. (**D**) Heatmap displaying log_2_ fold change (FC) of canonical NRF2 target genes after 24 and 72 h CDDO treatment, relative to vehicle-treated control. (**E-F**) ORA was performed on 72 h CDDO-treated AC16 samples through WEB-based Gene SeT AnaLysis Toolkit. The most upregulated (red) and downregulated (blue) significant (FDR<0.05) gene ontology (GO) biological processes and molecular functions (up to 10 pathways, ranked by enrichment score) are presented. Black points indicate -log_10_FDR. (**G**) Top 10 most upregulated KEGG pathways following ORA after 72 h CDDO treatment compared to vehicle-treated control samples, ranked by enrichment score. Black points indicate -log_10_FDR. (**H**) Ferroptosis KEGG pathway (hsa04216) with significantly upregulated DEGs after 72 h CDDO treatment relative to vehicle-treated control samples shaded in red according to log_2_FC.

Overrepresentation analysis (ORA) was subsequently performed on 72 h CDDO-treated DEGs. Gene ontology (GO) further corroborated the NRF2-specific nature of CDDO-mediated gene induction with the most highly enriched significant biological processes and molecular functions included cellular metabolic processes, cell redox homeostasis, response to xenobiotic stimulus (Fig. 10E), oxioreductase activity, antioxidant activity and electron transfer activity (Fig. 10F). GO cellular component analysis showed a significant decrease in collagen-containing extracellular matrix. Accordingly, the most downregulated GO terms also included extracellular matrix (ECM)-related terms (extracellular structure organisation, ECM structural component, ECM binding) and transforming growth factor beta (TGF-β) production (Fig. 10E-F). A notably downregulated GO biological process 72 h after CDDO treatment included the extrinsic apoptotic signalling pathway (Fig. 10E). ORA was also performed on upregulated DEGs using the Kyoto Encyclopedia of Genes and Genomes (KEGG) database. Further consistency with the NRF2-specific nature of CDDO was established whereby glutathione metabolism and metabolism of xenobiotics by cytochrome P450 were amongst the 10 most enriched KEGG pathways (Fig. 10G). Interestingly, ferroptosis was an enriched pathway and further analysis of this pathway highlighted 8 DEGs contributing to this result: *GCLC*, *GCLM*, *FTL*, *HMOX1*, *SLC40A1* (encoding ferroportin), *SLC7A11* and *FTH1*, highlighting strong correlation between glutathione homeostasis and ferroptotic pathways (Fig. 10H).

### 3.9 DOX promotes cell death mechanisms whilst inhibiting cell cycle in AC16 cells

Transcriptomic analysis on DOX treated AC16 cells (100 nM, 48 h) relative to vehicle-treated control cells resulted in the significant differential expression of >6000 genes of which 3934 were upregulated (Fig. 11A). ORA showed that amongst the most enriched biological processes were interleukin-4 production, TGF-β production and apoptotic processes involved in development (Fig. 11B). The top 10 most downregulated GO biological processes could all be attributed to cell cycle including kinetochore organisation and mitotic spindle assembly checkpoint (Fig. 11B). These themes of ECM and cell cycle perturbations were also observed following GO cellular component and molecular function analysis (Fig. 11C-D). ORA via the KEGG database highlighted taurine and hypotaurine metabolism as the most significantly enriched pathway, followed by p53 signalling pathway and ATP-binding cassette (ABC) transporters (Fig. 11E). Contrastingly, steroid biosynthesis and cell cycle were amongst the most downregulated KEGG pathways (Fig. 11E).

**Figure 11:**
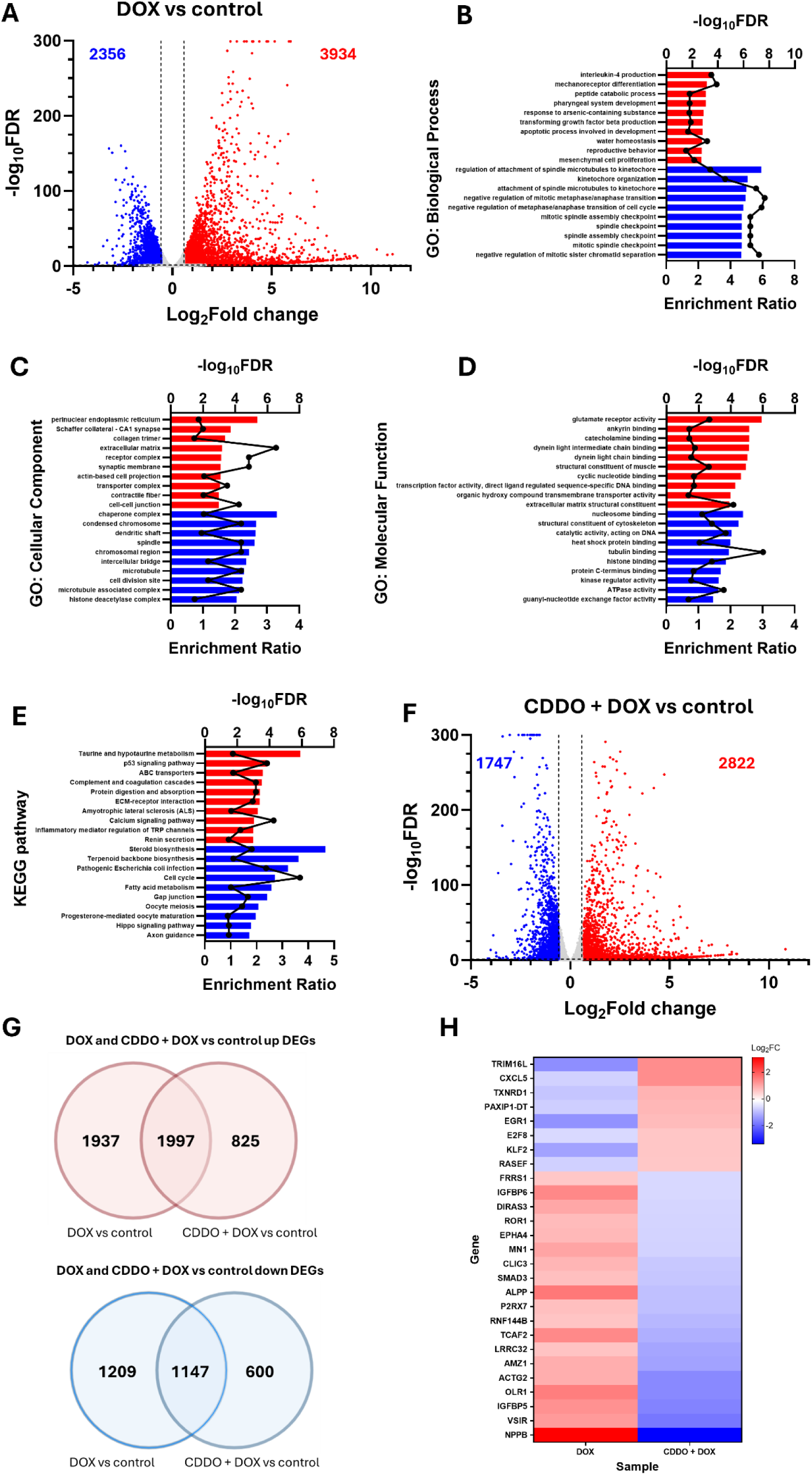
DOX has a large effect on the AC16 transcriptome. (**A**) Differential expression analysis volcano plots for 48 h, 100 nM DOX treatment compared to vehicle-treated control (n=3 biological repeats for all conditions). Blue and red points indicate downregulated and upregulated DEGs, respectively. (**B-D**) ORA was performed on 100 nM, 48 h DOX-treated AC16 samples through WEB-based Gene SeT AnaLysis Toolkit. The most upregulated (red) and downregulated (blue) significant (FDR <0.05) gene ontology (GO) biological processes, cellular components and molecular functions (up to 10 pathways, ranked by enrichment score) are presented. Black points indicate -log_10_FDR. (**E**) Top 10 most upregulated (red) and downregulated (blue) KEGG pathways following ORA after DOX treatment compared to vehicle-treated control samples, ranked by enrichment score. Black points indicate -log_10_FDR. (**F**) Differential expression analysis volcano plots for 24 h CDDO pre-treated (100 nM) AC16 cells followed by 48 h, 100 nM DOX treatment compared to vehicle-treated control (n=3 biological repeats for all conditions). Blue and red points indicate downregulated and upregulated DEGs, respectively. (**G**) Overlap of DOX-treated samples ± CDDO pre-treatment upregulated (red) or downregulated (blue) DEGs relative to vehicle-treated control in AC16 cells. (**H**) Heatmap depicting log_2_FC relative to control for DEGs with opposing directions of regulation between DOX and CDDO + DOX AC16 samples.

### 3.10 CDDO dampens DOX-mediated induction of apoptosis

Transcriptomic analysis of cells treated with DOX ± CDDO pre-incubation showed 4569 significant DEGs where 2822 were upregulated and 1747 downregulated (Fig. 11F).

Overlaying these data with DOX vs control DEGs showed a strong degree of overlap in these datasets with 1997 genes upregulated in both conditions and 1147 downregulated (Fig. 11G). However, a large number of genes were thus uniquely regulated by either CDDO + DOX or DOX alone in both directions relative to control samples (Fig. 11G). Interestingly, 27 genes had opposing directions of regulation between the datasets including *TRIM16L* and *TXNRD1* (Fig. 11H) which were notably amongst the most upregulated genes following 72 h CDDO treatment in isolation (Fig. 10B). KEGG analysis of these 27 genes did not yield any significant pathways; GO highlighted significance for cellular component insulin-like growth factor binding protein complex with insulin like growth factor binding protein 5/6 (IGFBP5/6) specifically implicated but no other significant terms (data not shown).

KEGG pathways identified via ORA from the DOX vs control dataset were explored in greater detail to identify the DEGs contributing to these pathways and how CDDO pre-treatment affected their regulation. In the context of the ABC transporters pathway, whilst CDDO + DOX vs control samples did not reverse the direction of activity, 25 of these genes including *ABCA13*, *ABCA1*, *ABCA12*, *ABCC11* and *ABCG1* were dampened in comparison to DOX vs control (Fig. 12A). Similarly, the downregulation of DEGs contributing to the inhibition of cell cycle within the DOX vs control dataset were largely diminished with CDDO pre-incubation, most notably with *SMAD3* where DOX treatment enriched its expression whereas this was significantly downregulated in the CDDO + DOX samples (Fig. 12B). The genes most upregulated following DOX treatment within the cell cycle KEGG pathway were *CDKN1A* (p21), *SFN* and *MDM2* whereas the most inhibited were *PLK1*, *CDKN2C* and *CCNB1* (Fig. 12B). KEGG pathway analysis of DOX treated samples relative to control samples also showed the p53 signalling pathway to be significantly enriched yet CDDO pre-treatment largely dampened these effects (Fig. 12C). Moreover, apoptosis was investigated as a specific pathway of interest considering the aforementioned effects of CDDO in the inhibition of extrinsic apoptosis (Fig. 10E) and that DOX has strong effects on cell viability (Fig. 3B). Key pro-apoptotic genes *BAX*, *CASP10* and *CASP3* were all significantly upregulated and *BCL2*, an anti-apoptotic gene, was significantly inhibited, indicating a pro-apoptotic phenotype in AC16 cells exposed to DOX (Fig. 12D). However, CDDO pre-treatment before DOX revealed no significant change in *BCL2* relative to control samples. Moreover, DOX-induced *FAS*, *APAF1* and *CASP10* increases relative to control cells were dampened with CDDO pre-treatment (Fig. 12D).

**Figure 12:**
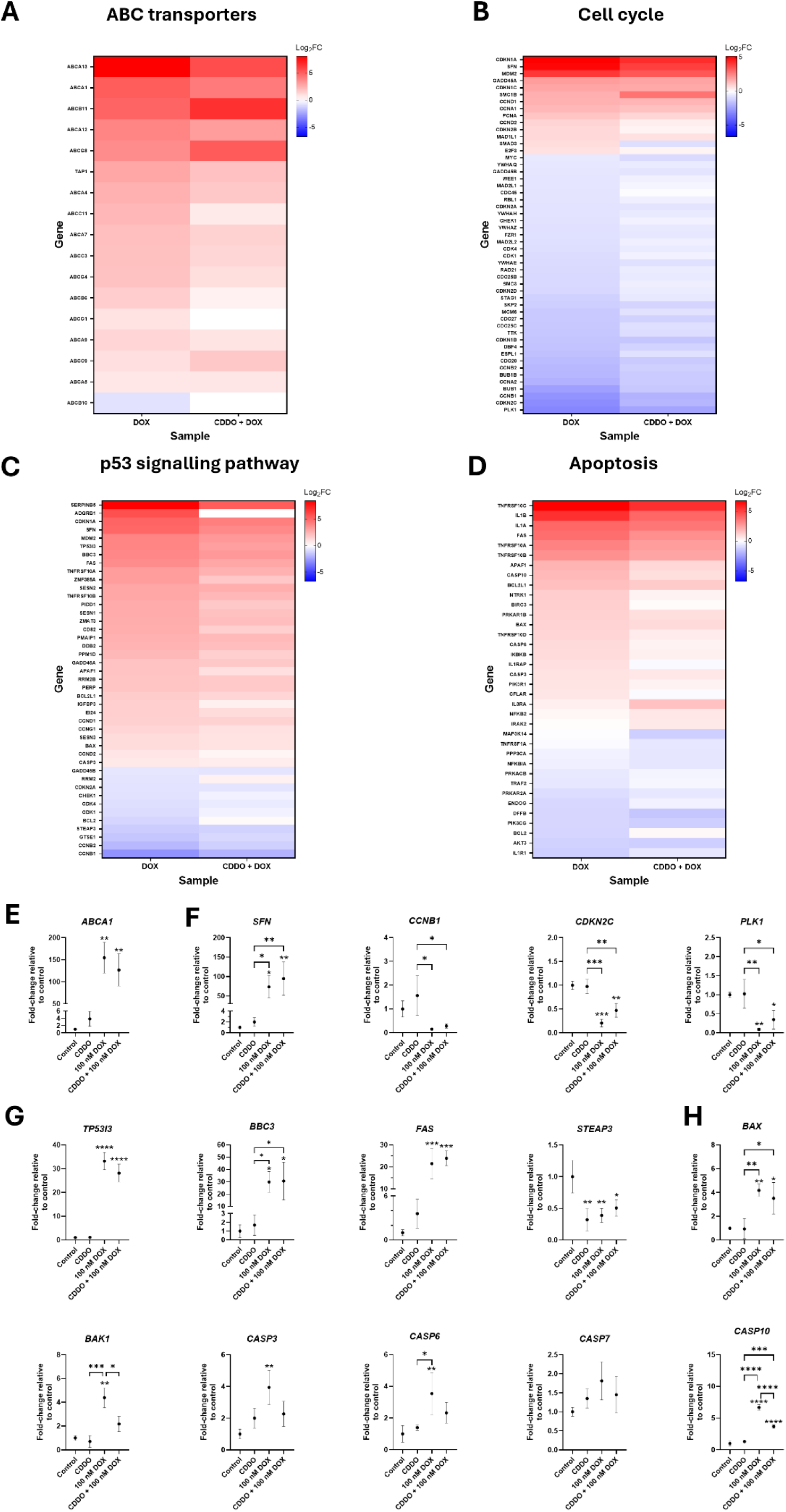
CDDO pre-incubation can dampen DOX-mediated effects on the AC16 transcriptome. (**A-B**) Heatmaps displaying log_2_FC gene expression data for DOX (100 nM, 48 h) treated AC16 cells ± 24 h pre-treatment with 100 nM CDDO compared to vehicle-treated control samples within the KEGG pathways ATP-binding cassette (ABC) transporters (hsa02010) and cell cycle (hsa04110). (**C-D**) Heatmaps displaying log_2_FC gene expression data for DOX (± pre-treatment with CDDO)-treated AC16 cells compared to vehicle-treated control samples within the KEGG pathways p53 signalling pathway (hsa04115) and apoptosis (hsa04210). (**E-F**) qPCR analysis in AC16 cells vehicle-treated (control), CDDO-treated (72 h, 100 nM), DOX-treated (100 nM, 48 h) or CDDO + DOX-treated (100 nM 24 h CDDO pre-treatment, 100 nM, 48 h DOX treatment) for selected genes within the ABC transporters (E), cell cycle (F), p53 signalling (G) and apoptosis pathways (H). qPCR data are normalised to control and presented as mean ± SD (n=3). Asterisks above data points indicate significant difference to control whereas asterisks above brackets indicate significant difference between pairwise comparisons following a one-way ANOVA with Tukey’s post-hoc analyses (*p<0.05, **p<0.01, ***p<0.001, ****p<0.0001).

Re-confirmation of the effects of DOX and CDDO + DOX were performed on *ABCA1* (Fig. 12E), *SFN*, *CCNB1*, *CDKN2C* and *PLK1* (Fig. 12F). *ABCA1*, *SFN* and *CCNB1* were all significantly enriched after DOX treatment relative to control samples whereas *CDKN2C* and *PLK1* were inhibited at the transcript level by DOX (Fig. 12E-F). Whilst not restored to baseline, CDDO + DOX treatment less significantly inhibited *CDKN2C* and *PLK1* expression (Fig. 12F). Re-confirmation of p53 signalling and apoptosis KEGG pathways was undertaken via qPCR using GAPDH as a housekeeping control which was stable under DOX treatment (Fig. S6). DOX significantly induced *TP53I3*, *BBC3*, *FAS* and significantly inhibited *STEAP3* and CDDO was not effective for diminishing these effects (Fig. 12G). As seen in the RNA-seq analyses, *BAX*, *BAK1*, *CASP3*, *CASP6*, and *CASP10* were significantly upregulated upon DOX treatment; *CASP7* was unaffected (Fig. 12H). Notably, CDDO pre-incubation significantly reduced the effects of DOX for these genes (Fig. 12H).

The effects of DOX on NRF2-related transcriptional activity were also evaluated within the RNA-seq datasets. Many canonical NRF2 targets within the aforementioned manually curated gene array (Fig. 10D) were downregulated with DOX treatment such as *HKDC1*, *NQO1* and *TXNRD1* (Fig. S7). Interestingly, *NFE2L2* was downregulated by 0.53 log_2_ fold change and *KEAP1* expression was also significantly ablated by 0.62 log_2_ fold change (Fig. S7). Pre-treatment with CDDO for 24 h stabilised the expression of NRF2 targets including *NQO1*, *TXNRD1* and *GSR* (Fig. S7).

## 4 Discussion

In this study, we investigated the contribution of CDDO-mediated NRF2 activation towards the alleviation of DOX-induced cytotoxicity in human cardiomyocytes. Whilst the exact mechanisms through which DOX exerts these effects are multifactorial, oxidative stress and mitochondrial dysfunction are well documented hallmarks of its adverse events [46,47], leading to cardiomyocyte cell death, the predominant and non-replenishable cardiac cell type [48], and subsequent tissue dysfunction. As a master regulator of antioxidant responses alongside many other pro-survival mechanisms, including detoxification, anti-inflammation, anti-apoptosis and autophagy regulation [49], NRF2 has recently emerged as a candidate target in the alleviation of anthracycline-induced cardiotoxicity [50]. However, the exact mechanisms through which NRF2 exerts cytoprotective effects in cardiomyocytes has not been conclusively evidenced. Moreover, the selection of appropriate pharmacological molecules which limit off-target effects is critical in the context of doxorubicin cardiotoxicity, given its clinical utility as an antineoplastic agent and the dual functions of NRF2 in cancer [51].

We selected the AC16 cell line, a fusion between primary cells from human ventricular cardiac tissue and SV40 transformed human fibroblasts [52]. This cell line provided a valuable *in vitro* model of DOX-induced cytotoxicity in a clinically-relevant cell type with amenable application of biomolecular tools to study NRF2 dynamics. NRF2 induction was achieved via incubation of AC16 cells with CDDO, a molecule with considerable clinical trial history in the context of cancer, chronic kidney disease, COVID-19, hypertension and neurological disorders [53]. Interestingly, a CDDO-analogue, omaveloxolone, recently received FDA-approval in Friedreich’s ataxia [54]. CDDO successfully induced NRF2 protein expression and downstream activity as a transcription factor, driving the expression of well-reported cytoprotective and canonical targets TXNRD1, NQO1, FTH1 and GCLM at both the RNA and protein level (Fig. 1). Moreover, in single-cell analyses using lentiviral overexpression of fluorescently-labelled NRF2, rapid induction of NRF2-mNG was measured; 1.25-fold after 10 min and 1.6-fold by 30 min post-CDDO treatment (Fig. 2C). This is in line with other NRF2 stabilisers [55–57]. Fluorescence correlation spectroscopy (FCS) was utilised to quantify significant increases in both cytoplasmic and nuclear NRF2-mNG signals (Fig. 2B). Alternative quantitative single-cell models of NRF2 dynamics have been reported but could not determine exact protein molecule counts [58,59]. This highlights an advantage of FCS as a technique to study NRF2 activators and perhaps in compound screening [60].

AC16 cells were incubated with CDDO for 24 h prior to DOX exposure to allow the upregulation of downstream NRF2 targets. This resulted in significant protection of cells against 100 nM and 1 µM DOX (Fig. 3). Interestingly, peak DOX levels in patient plasma after intravenous delivery is ∼1 µM, rapidly decreasing to 30-75 nM [61–63]. As such, 100 nM and 1 µM could be considered a clinically-relevant concentration of DOX. Genetic upregulation of NRF2 cytoprotection against DOX was further explored using the aforementioned NRF2-mNG overexpression AC16 cells, which were significantly more resilient with regards to cell viability than WT AC16 cells (Fig. 4F-G). Moreover, the specific contribution of NRF2 to CDDO-mediated effects was evaluated using siRNA inhibition of *NFE2L2* or *KEAP1* which ameliorated any protection against DOX toxicity; this model also supports the hypothesis that NRF2 promotes AC16 survival in the presence of 100 nM DOX whereby *KEAP1* downregulation demonstrated significantly greater cell viability than *NFE2L2* suppression (Fig. 5G). CDDO effects on the AC16 transcriptome were explored via RNA-seq. The number of DEGs activated or inhibited after 24 h and 72 h CDDO treatment were relatively low (≤147; Fig. 10A-C) but were largely specific to the NRF2 pathway either through known, canonical downstream targets within a manually curated list (Fig. 10D) or via activation of well-characterised pathways such as glutathione metabolism or xenobiotic detoxification (Fig. 10E-F). These data corroborate the recent evidence that CDDO may act upon a single, highly reactive cysteine residue of KEAP1; C151 to exert highly-specific NRF2 activation, in contrast to other agonists such as CDDO-Im or sulforaphane which modulate multiple KEAP1 cysteine residues [64].

Despite clinical utility for >60 years, the specific mechanisms of DOX-induced cardiotoxicity are yet to be fully explored. Data presented in this study facilitate the bridging of this knowledge gap via transcriptomic analysis of DOX-treated human cardiomyocytes at clinically-relevant doses. Pathways and GO terms associated with cell cycle were significantly downregulated in DOX-treated AC16 cells relative to control cells. DEGs pertaining to this effect included the significant upregulation of genes encoding established cell cycle arrest proteins CDKN1A (p21), SFN and MDM2 [65–67] with subsequent inhibition of PLK1 and CCNB1, both key drivers of oncogenic mitosis [68,69]. Several known DOX-upregulated mechanisms emerged as the most enriched pathways in this RNA-seq dataset such as the p53 signalling pathway, ABC transporters (likely a feedback process to promote DOX efflux from cells [70]) and apoptosis [71] (Fig. 11). Apoptosis is one of many cardiomyocyte programmed cell death pathways that provide a significant contribution to cardiovascular toxicity induced by DOX [72,73] yet the specific mechanism for cell death may be dependent upon the DOX concentration. In a study on acute myeloid leukaemia, a disease in which DOX in a frontline therapy, high doses of 5 µM DOX induced necrosis whereas more clinically-relevant doses (0.5-1 µM) similar to those utilised in this study imparted late apoptotic cell death [74].

Intriguingly, DOX also enriched processes surrounding ECM dysregulation for example the GO terms collagen trimer, extracellular matrix and extracellular matrix structural component were upregulated (Fig. 11C-D). Fibrosis is a contributing factor to DOX-induced cardiotoxicity, mainly driven by the activation of inflammatory responses and cardiomyocyte cell death, a non-regenerative cell population [75]. Lost cells are thus replenished largely via immune cell infiltration and fibroblasts [76] which secrete ECM components and increase cardiac stiffness which promotes many of the DOX-induced physiological defects on cardiac function leading to impaired contractility and diminished ventricular function [77,78]. Notably, one of the major effects of CDDO, uncovered via the transcriptomic data presented here, suggests an anti-fibrotic phenotype is produced with inhibition of TGF-β production, response and reduced ECM binding (Fig. 10). This may highlight a major process through which CDDO exerts cardioprotective effects; NRF2 activation is also known to dampen cardiac fibroblast activation and fibrotic effects of TGF-β *in vitro* and *in vivo* [79,80]. Results from the RNA-seq analysis must be validated further in subsequent studies but indicate areas of interest and support the understanding regarding the impact of DOX and NRF2 cytoprotection within a cardiac context. A recent and comprehensive study into temporal and concentration-dependent DOX cardiotoxicity corroborate the data presented here whereby DOX promotes apoptosis (perhaps linked to mitochondrial damage) and fibrosis in AC16 cells [81].

Given the potential for excessive NRF2 activation to promote tumour resistance, we assessed CDDO effects on multiple human cancer cell lines from lung adenocarcinoma and pancreatic lineages. WT A549 cells, with constitutively active NRF2 thus modelling human lung cancer which demonstrate frequent mutations (∼28%) within the NRF2/KEAP1 pathway [31,82], was relatively resistant to DOX treatment but was significantly sensitised by NRF2 knockout (Fig. S4A). This may suggest that DOX would be more efficacious against tumours of the lung which have typical NRF2 signalling with low baseline activity and yet should be relatively ineffective against high NRF2-expressing tumours. Furthermore, patients with low-NRF2 malignancies could be treated with lower doses of DOX, thus aiding the alleviation of off-target cardiotoxic events. This patient stratification according to tumour genotype provides an avenue for personalised medicine approaches. In contrast to AC16 cells, CDDO had no beneficial impact on A549 cells with WT or NRF2^-/-^ genotype and may even have promoted its cytotoxic effects (Fig. 8B-C). Cell lines from pancreatic cancer lineage were also selected due to previous evidence supporting a role of NRF2 in chemotherapeutic and radiotherapeutic resistance [42]. Whilst this previous report linked the SUIT-2 cell line with greater baseline NRF2 expression (driven by reduced KEAP1 protein) and glutathione activity than MIA PaCa2, in our study we could not determine significant difference in NRF2 or KEAP1 protein levels (Fig. 9 and Fig. S5). Functionally, both SUIT-2 and MIA PaCa2 cells were relatively resistant to DOX treatment and CDDO did not promote cell viability, concurring the A549 results, despite increases in NRF2 protein expression and downstream activity measured via TXNRD1 and GCLM expression (Fig. 9). This small proof-of-concept study provides an intriguing prospect whereby CDDO may promote AC16 survival upon exposure to DOX, in part via NRF2 and yet the same upregulation of NRF2 in cancer cells may not impart resistance. We highlight this hypothesis as a key area for future research which would require more detailed and in-depth analysis.

The limitation of this study that should be acknowledged is the utilisation of the AC16 cell line which, whilst derived from human cardiomyocytes and fibroblasts, were selected to provide molecular understanding about NRF2-mediated protection against DOX cytotoxicity within a cardiac context. For greater translational research for replicating cardiovascular biology, *in vivo* models should be used in further study or *in vitro* models which capture characteristics of the human cardiac tissue such as induced pluripotent stem cells or 3D spheroid cultures [25]. For example, the dividing nature of AC16 cells through fusion with SV40-transformed fibroblasts is not replicative of mature cardiomyocytes which do not proliferate. Whilst results here have highlighted the effects of DOX on cell cycle and ABC transporters, we appreciate that these pathways would not be as relevant specifically in post-mitotic cardiac cells. Despite this, they are prominent and notable results within this study. Moreover, AC16 cells proved advantageous when exploring NRF2 dynamics through the use of multiple experimental approaches which would have been challenging to implement *in vivo* or other *in vitro* models. Despite these limitations, this study provides novel insights into DOX cardiotoxicity and provides substantial evidence that NRF2 activation may provide a route for its alleviation.

## 5 Conclusions

Overall, this study provides valuable insight into the mechanisms of DOX-induced cytotoxicity in human cardiac cells. Activation of NRF2 has been highlighted as a robust and effective mitigation strategy for the alleviation of DOX-mediated cardiotoxicity which can be potently and specifically targeted via CDDO therapy. Novel insights into the mechanisms through which CDDO and NRF2 exert this protection such as via antioxidant, anti-ferroptotic and anti-fibrotic enrichment have been presented, whilst maintaining the antitumorigenic effects of the chemotherapeutic. Importantly, CDDO poses excellent clinical translation potential therefore the insights provided from this study could have real impact for patients undergoing chemotherapy.

## 6 Supplementary materials

**Figure S1:**
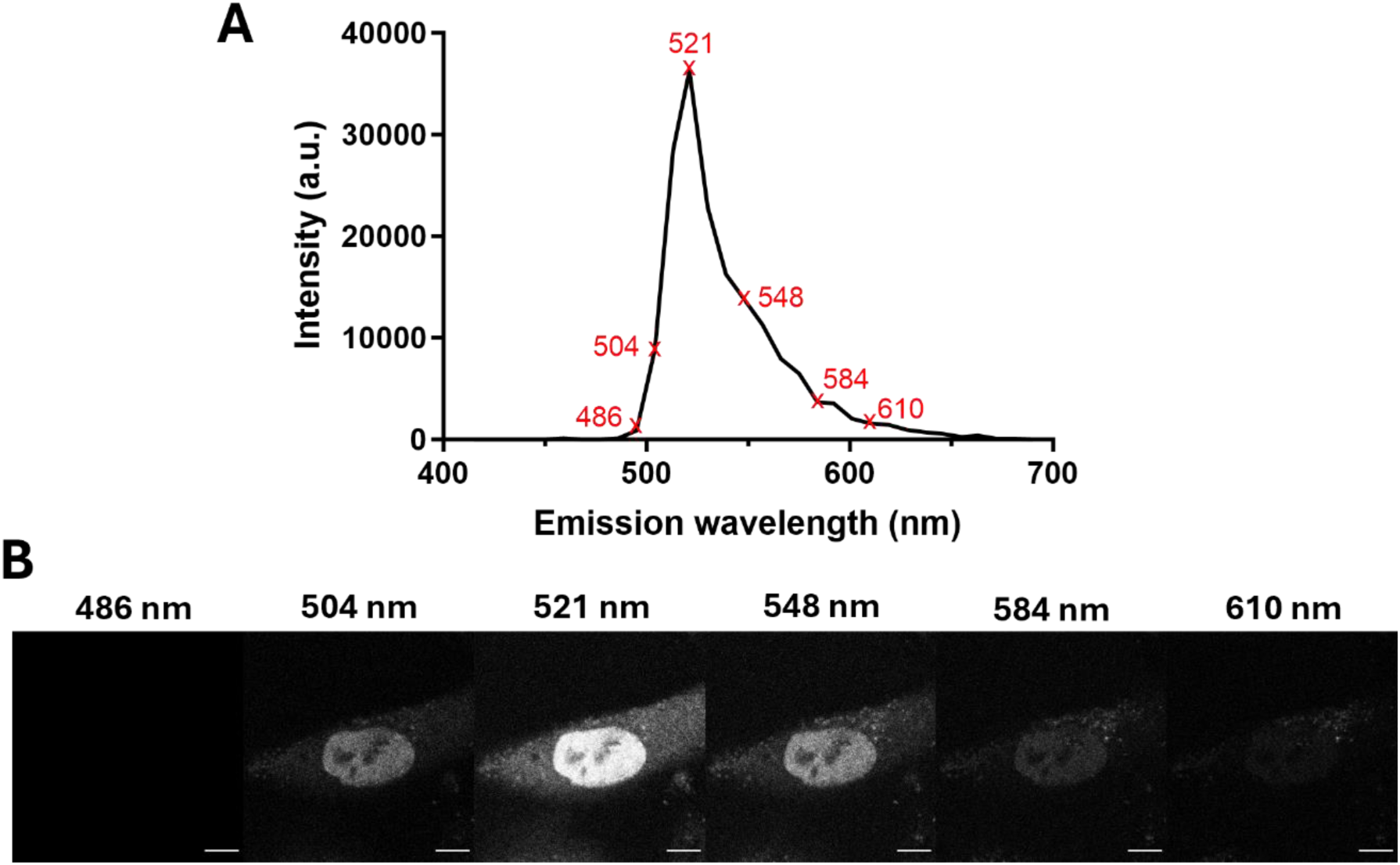
Lambda scan of NRF2-mNG-expressing AC16 cells. (**A**) Emission spectral profile following excitation at 488 nm expressed in arbitrary units (a.u.). Red crosshairs indicate emission wavelengths for which representative images (**B**) are provided. Scale bar is 10 µm.

**Figure S2:**
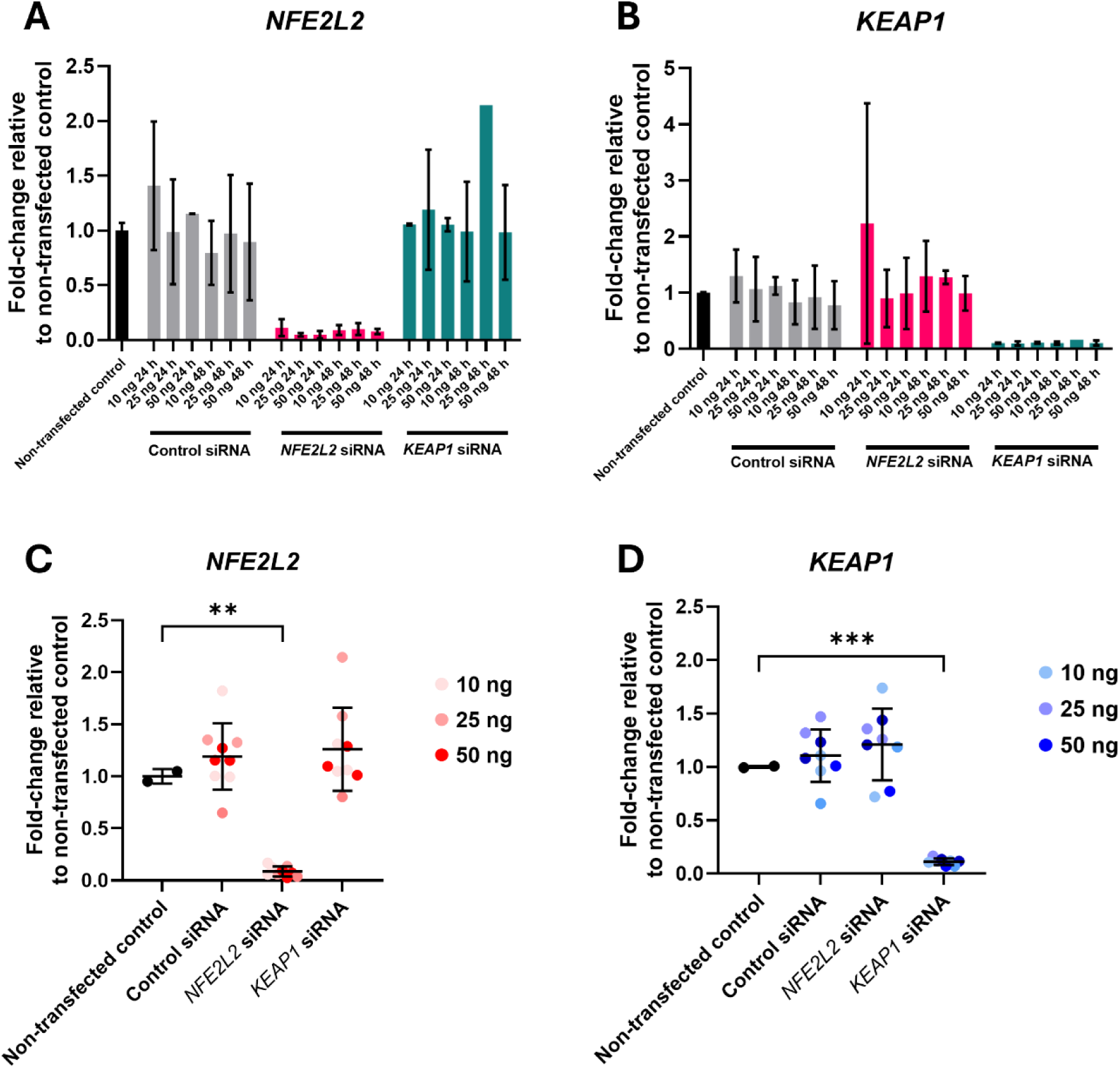
Optimisation of siRNA transfections in AC16 cells measured at the transcript level. *NFE2L2*, *KEAP1* or control siRNA were transfected into AC16 cells at 3 concentrations (10, 25 or 50 nM) and RNA harvested 24 or 48 h later. The mRNA expression of (**A**) *NFE2L2* or (**B**) *KEAP1* was subsequently measured by qPCR and normalised to expression within non-transfected cells. Data are mean ± SD (n=2 per condition). Pooled qPCR data across all concentrations and time points was analysed to determine siRNA efficacy against (**C**) *NFE2L2* and (**D**) *KEAP1*; data are mean ± SD (n≥7). Statistical significance was evaluated using a one-way ANOVA with Tukey’s correction. **p<0.01, *** p < 0.001.

**Figure S3:**
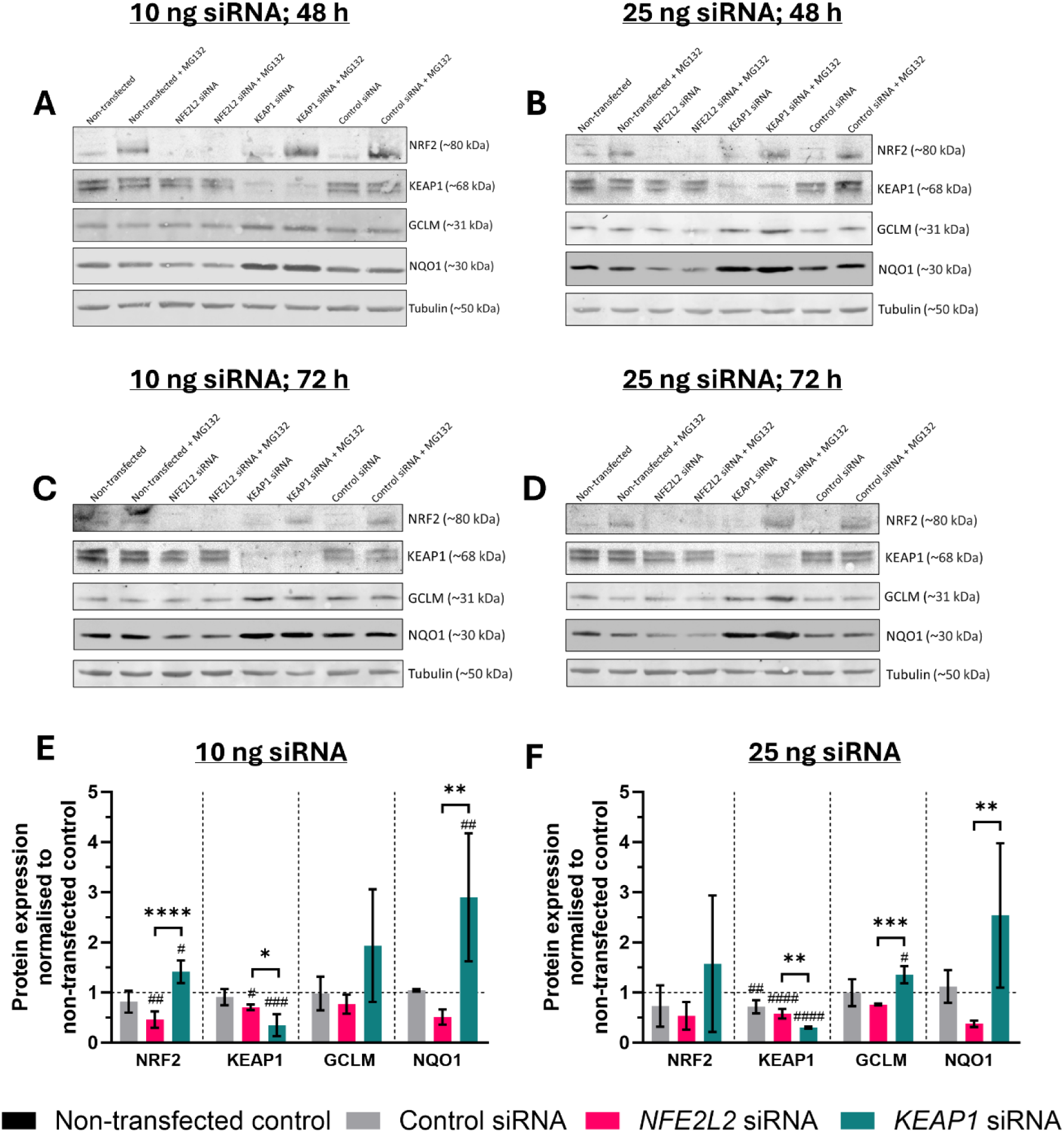
Optimisation of siRNA transfections in AC16 cells measured at the protein level. (**A-D**) Representative immunoblots of AC16 cells following transfection with (A) 10 ng siRNA, 48 h; (B) 25 ng siRNA, 48 h; (C) 10 ng siRNA, 72 h; (D) 25 ng siRNA, 72 h. Cells were treated ± 10 µM MG132 (2 h). (**E-F**) Basal, untreated protein quantification of NRF2, KEAP1, GCLM and NQO1 after (E) 10 ng and (F) 25 ng siRNA, respectively; pooled data from 48 and 72 h. Data are normalised to non-transfected, untreated AC16 cells (represented as dotted line) and presented as mean ± SD (n=4). Statistical significance was determined via one-way ANOVAs with Tukey’s post-hoc analyses. #p<0.05, ##p<0.01, ###p<0.001, ####p<0.0001 relative to non-transfected controls. *p<0.05, **p<0.01, ***p<0.001, ****p<0.0001 between groups.

**Figure S4:**
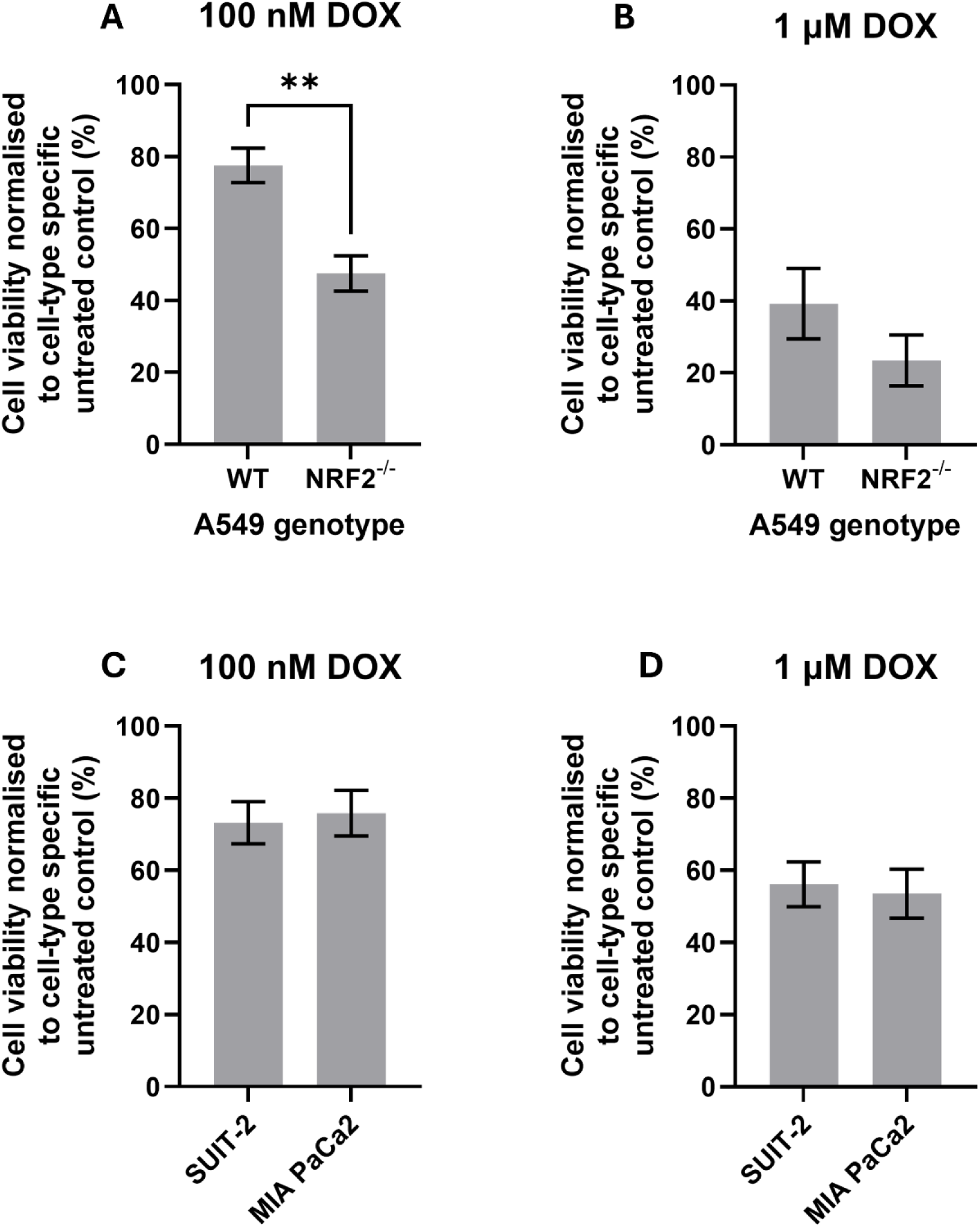
NRF2^-/-^ A549 cells are more sensitive to DOX than WT A549 cells whereas pancreatic cancer cells are comparable. (**A-B**) Cell viability of WT and NRF2^-/-^ A549 cells in response to 48 h treatment with (A) 100 nM or (B) 1 µM DOX, respectively. (**C-D**) Cell viability of SUIT-2 and MIA PaCa2 cells in response to 48 h treatment with (C) 100 nM or (D) 1 µM DOX, respectively. Data are normalised to genotype or cell-type specific untreated controls and reported as mean ± SEM (n=3). Statistical significance was determined using an unpaired, two-tailed Student’s t-test. **p<0.01.

**Figure S5:**
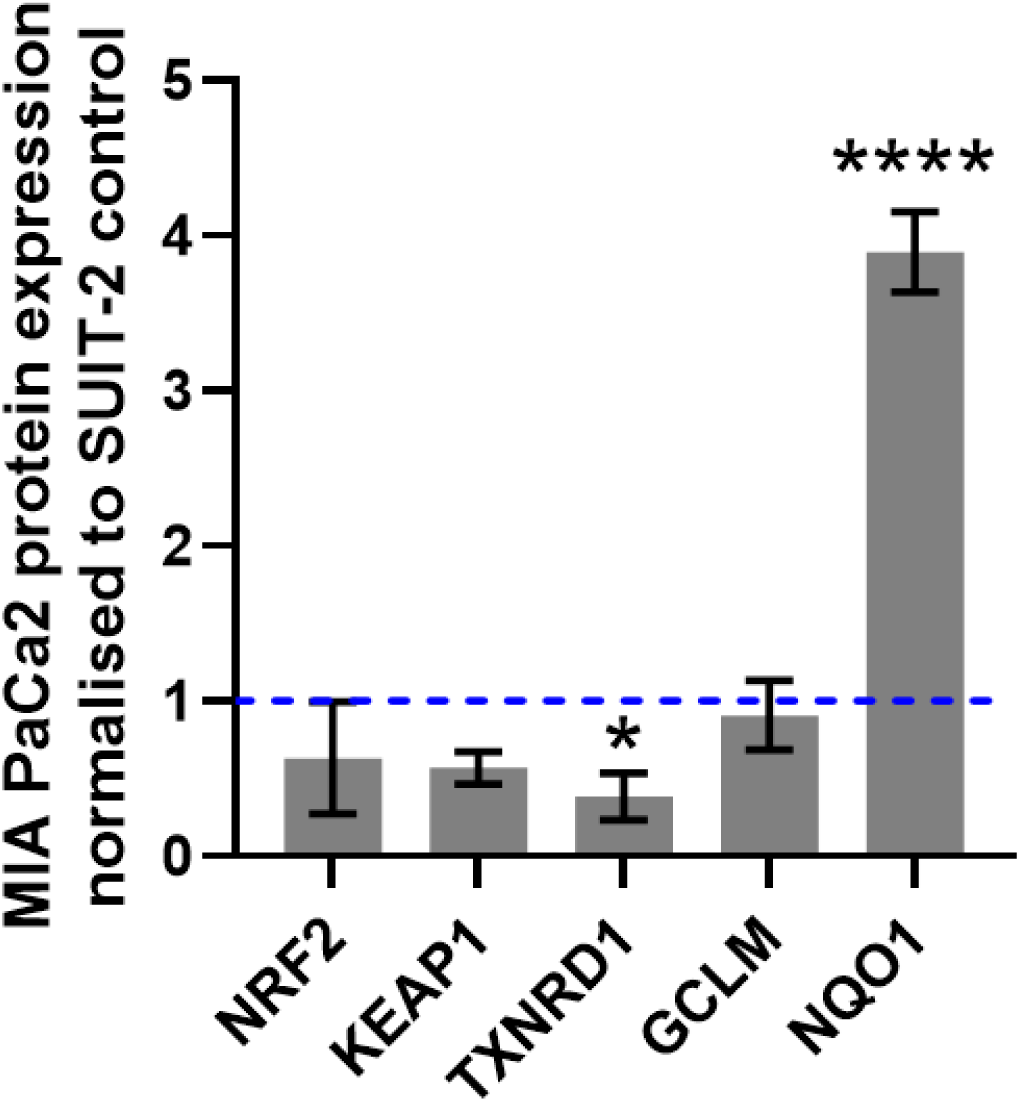
SUIT-2 and MIA PaCa2 have similar NRF2 expression. MIA PaCa2 protein quantification following Western blot analysis normalised to SUIT-2 (blue line) protein expression in untreated cells. Data are mean ± SD (n=3). Statistical significance was determined via a one-way ANOVA with Tukey’s post-hoc analysis. *p<0.05, ****p<0.0001 relative to SUIT-2.

**Figure S6:**
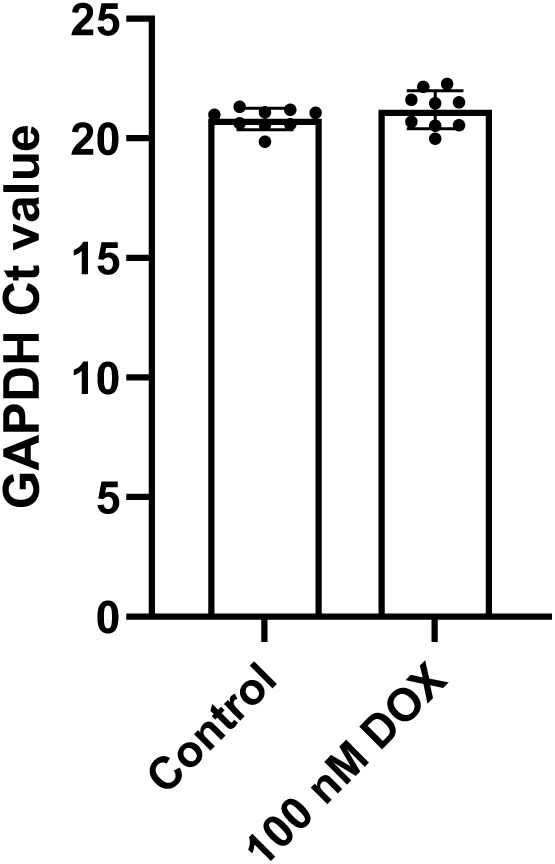
Ct values for GAPDH gene expression in AC16 cells under control (untreated) and 100 nM DOX (48 h incubation) backgrounds. Data are mean ± SD (n=8). No significant differences observed following an unpaired Student’s t-test.

**Figure S7:**
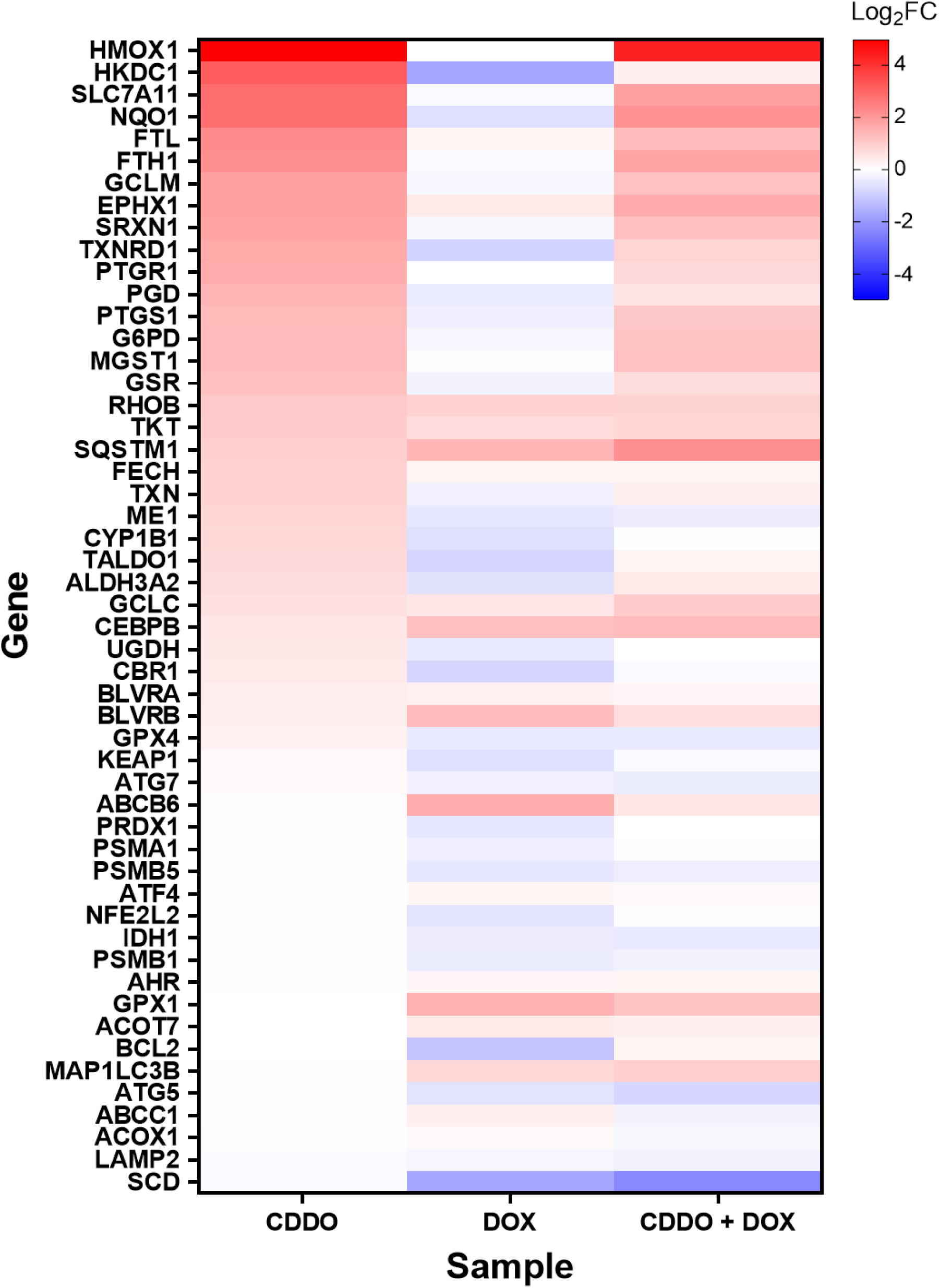
CDDO prevents DOX-mediated inhibition of NRF2 signalling pathways. All analyses were performed relative to control, vehicle-treated samples. CDDO indicates 72 h exposure to 100 nM CDDO, DOX indicates 48 h 100 nM DOX, CDDO + DOX indicates 24 h pre-treatment with 100 nM CDDO prior to 48 h 100 nM DOX.

**Table S1:**
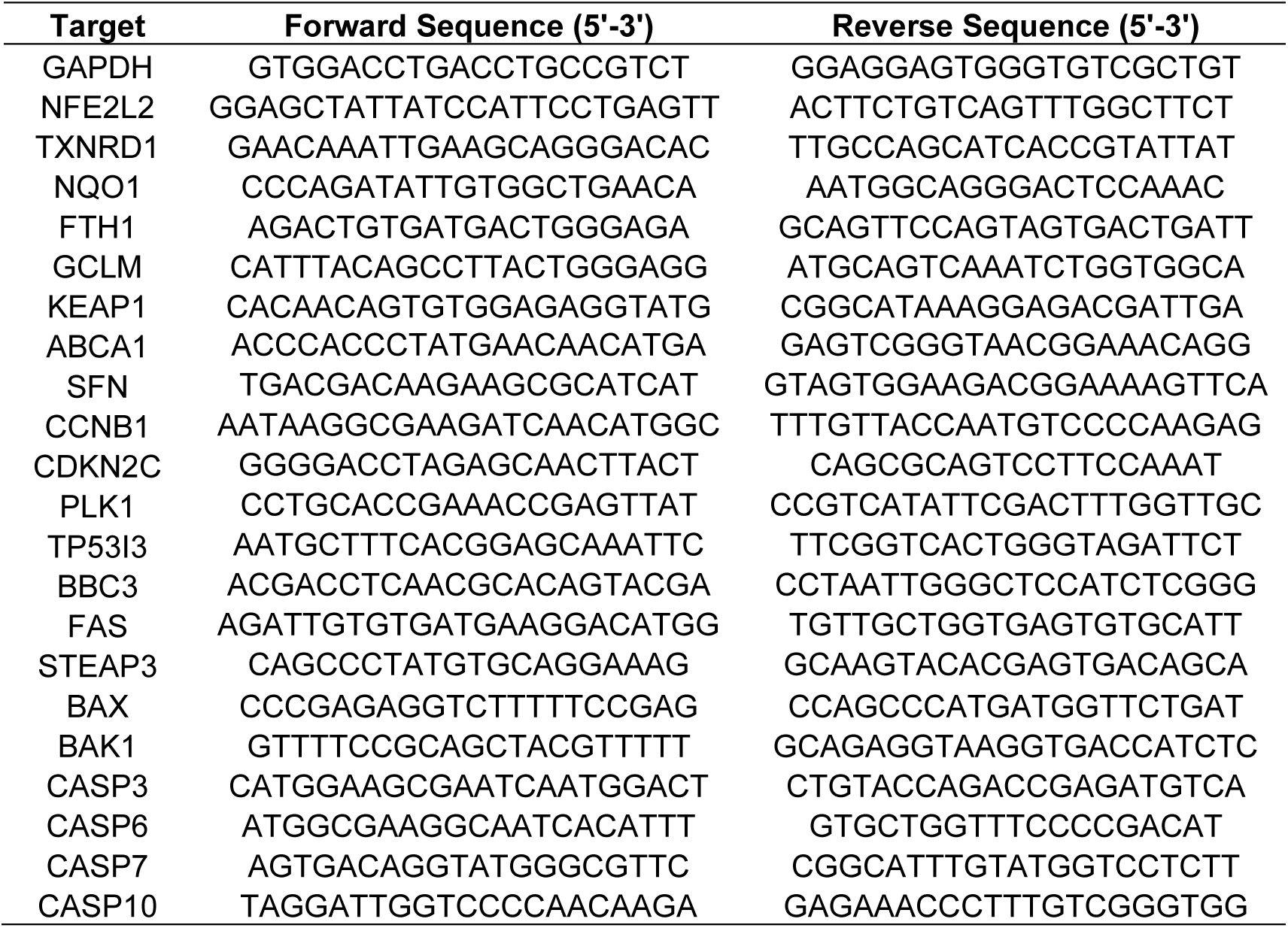
Primers utilised for qPCR experiments.

**Table S2:** Manually curated consensus NRF2 target genes.

|  |  |  |
| --- | --- | --- |
| ABCB6 | FTH1 | PGD |
| ABCC1 | FTL | PRDX1 |
| ACOT7 | G6PD | PSMA1 |
| ACOX1 | GCLC | PSMB1 |
| AHR | GCLM | PSMB5 |
| ALDH3A2 | GPX1 | PTGR1 |
| ATF4 | GPX4 | PTGS1 |
| ATG5 | GSR | RHOB |
| ATG7 | HKDC1 | SCD |
| BCL2 | HMOX1 | SLC7A11 |
| BLVRA | IDH1 | SQSTM1 |
| BLVRB | KEAP1 | SRXN1 |
| CBR1 | LAMP2 | TALDO1 |
| CEBPB | MAP1LC3B | TKT |
| CYP1B1 | ME1 | TXN |
| CYP1B1 | MGST1 | TXNRD1 |
| EPHX1 | NFE2L2 | UGDH |
| FECH | NQO1 |  |

## Data availability

RNA-seq data are deposited at the Gene Expression Omnibus [83]; GEO ID will be uploaded upon acceptance. Other data are available upon request.

## CRediT Author Contribution

**James A. Roberts:** Conceptualisation, methodology, validation, formal analysis, investigation, data curation, writing – original draft, visualisation, project administration. **Michael Batie:** methodology, software, formal analysis, resources, data curation, visualisation, writing – review and editing. **Amy H. Ponsford:** methodology, investigation, resources, writing – review and editing, supervision, project administration. **Jonathan Poh:** methodology, investigation, resources, writing – review and editing. **Benjamin J. Hewitt:** Writing – review and editing. **Hannah F. Botfield:** Writing - review and editing. **Lisa J. Hill:** Writing – review and editing. **Christopher M. Sanderson:** Conceptualisation, methodology, resources, supervision, project administration, funding acquisition. **Sonia Rocha:** Conceptualisation, methodology, resources, writing – review and editing, supervision, project administration. **Parveen Sharma:** Conceptualisation, methodology, validation, resources, writing – review and editing, supervision, project administration, funding acquisition.

## Acknowledgements

The authors would like to acknowledge the Centre for Cell Imaging at the University of Liverpool for imaging support. Graphical abstract was produced using BioRender.

